# Native proline-rich motifs exploit sequence context to target actin-remodeling Ena/VASP proteins

**DOI:** 10.1101/2021.03.22.436451

**Authors:** Theresa Hwang, Robert A. Grant, Meucci W. Ilunga, Venkatesh Sivaraman, Amy E. Keating

## Abstract

The human proteome is replete with short linear motifs (SLiMs) of 4-6 residues that are critical for protein-protein interactions, yet the importance of the sequence surrounding such motifs is underexplored. We devised a proteomic screen to systematically examine the influence of SLiM sequence context on protein-protein interactions. Focusing on the EVH1 domain of ENAH, an actin regulator that is upregulated in invasive cancers, we screened 36-residue proteome-derived peptides for binding. We discovered a pocket on the ENAH EVH1 domain that diverged from its orthologs to recognize extended SLiMs, and we found that proteins with two EVH1-binding SLiMs can wrap around a single domain. We also found that the ciliary protein PCARE uses an extended 23-residue region to obtain higher affinity than any known ENAH EVH1-binding motif. Our screen provides a way to uncover the effects of broader proteomic context on motif-mediated interactions, revealing diverse mechanisms of contextual control over EVH1 interactions and establishing that SLiMs can’t be fully understood outside of their native context.

## Introduction

Interactions between modular interaction domains and short linear motifs (SLiMs) direct a broad range of intracellular functions, from protein trafficking to substrate targeting for post-translational modifications. To faithfully propagate signals, SLiMs must recognize the correct interaction partners within the cellular environment. But how interaction specificity is achieved is enigmatic. SLiMs, which occur as 3-10 consecutive amino acids in intrinsically disordered regions of proteins are degenerate and have low complexity, meaning they are defined by just a few key residues or motif features. Crystal structures of SH3, WW, and PDZ domains bound to SLiMs typically reveal 3-6 residues docked into a shallow groove (Lim et al., 1994; Macias et al., 1996; Schultz et al., 1998). In addition, the expansion of modular interaction domain families in metazoan proteomes has led to hundreds of domains that share overlapping SLiM-binding specificity profiles yet carry out distinct functions in the cell (Bhattacharyya et al., 2006). How high-fidelity interactions are maintained between low complexity SLiMs and cognate recognition domains remains poorly understood for many pathways.

Most SLiM research has centered around defining the “core SLiM,” or the minimal set of amino acids sufficient to bind to a given domain. High-throughput approaches, such as peptide phage display using libraries of 7-16-residue peptides, have been instrumental for advancing our understanding. But these assays don’t probe how the sequences surrounding core SLiMs affects their interactions (Ivarsson et al., 2014; Teyra et al., 2017; Tonikian et al., 2008; Davey et al., 2017;). There is increasing evidence that surrounding sequence critically influences SLiM interaction affinity and specificity (Palopoli et al., 2018; Prestel et al., 2019). For example, an alpha helical extension C-terminal to a SLiM in ankyrin-G enables high affinity (K_D_ = 2.6 nM) and selective interactions with the GABARAP subfamily of Atg8 proteins, by making additional contacts with the GABARAP interface (Li et al., 2018a). In addition, the presence of aromatic residues directly flanking a SLiM in Drebrin prevents its interaction with Homer (Li et al., 2019), demonstrating that SLiM sequence context can also disfavor proteinprotein interactions.

The actin interactome contains many proline-rich SLiMs and many proline-binding modules such as SH3, WW, and EVH1 domains that participate in regulating actin dynamics (Holt and Koffer, 2001). Although the extent to which these domains crossreact or bind selectively in the cell is unknown, sequence elements surrounding linear, proline-rich motifs could play an essential role in directing specific interactions. Therefore, we sought to uncover the impact of sequence context on SLiM-mediated interactions with the EVH1 domain of the actin-regulating Ena/VASP proteins.

Ena/VASP proteins form a family of cytoskeletal remodeling factors that are recruited to different regions of the cell by binding proline-rich SLiMs via their N-terminal EVH1 domains and promoting actin polymerization via their C-terminal EVH2 domains. The family is implicated in many cellular functions such as axon guidance and cell adhesion (McConnell et al., 2016; Scott et al., 2006). The Ena/VASP EVH1 domain recognizes the SLiM [FWYL]PXΦP, where X is any amino acid and Φ is any hydrophobic residue (Ball et al., 2000). This motif, referred to in this paper as the *FP4 motif*, adopts a polyproline type II (PPII) helix structure and binds weakly to the EVH1 domain (Prehoda et al., 1999). Searching for this core FP4 motif in the human proteome yields 4994 instances. This number of potential interaction partners is very large, and although spatial, structural, and temporal context impose additional determinants for cellular interaction (Bugge et al., 2020), the abundant motif matches raise the question of whether sequence elements beyond the FP4 SLiM affect molecular recognition.

We used a new screening approach to uncover examples of how the sequence context surrounding the core Ena/VASP FP4 SLiM affects binding specificity in the proteome. Our unbiased screening method, MassTitr, identified 36-residue human proteome-derived peptides that bind to the ENAH EVH1 domain with a range of affinities. To our knowledge, this is the first use of a high-throughput screening method to systematically discover and characterize both local and distal sequence elements that impact SLiMs. By analyzing features of high-affinity binders, we identified distinct ways in which sequence elements surrounding proteomic FP4 SLiMs impact binding affinity and specificity to ENAH. Our work provides insight into how selective interactions are maintained in proline-rich motif-mediated signaling networks and highlights the importance of considering sequence context when investigating SLiM-mediated interactions. Our pipeline serves as a blueprint to map and predict how sequence context surrounding SLiMs impact protein-protein interactions on a proteome-wide scale.

## Results

### MassTitr identifies ENAH EVH1 long domain-binding peptides from the human proteome

To identify ENAH EVH1 binders in the human proteome, we applied a new screen called MassTitr. MassTitr is a SORT-SEQ method that is based on fluorescence-activated cell sorting (FACS) of a library of peptide-displaying bacteria and subsequent deconvolution of signals by deep sequencing. As shown in Figures 1A and B, peptide-displaying *Escherichia coli* cells are sorted into bins according to their binding signals across a range of protein concentrations, and the binding signal for each peptide at each protein concentration is extracted by deep sequencing each bin. Two advantages of this method over phage display using the p8 gene are that MassTitr supports screening of long peptides, and it leads to identification of binders with a broad range of affinities. MassTitr is similar in concept to the yeast-based Tite-seq, which Adams et al. applied to study CDR loops of engineered anti-fluorescein scFvs (Adams et al., 2016).

**Figure 1.**
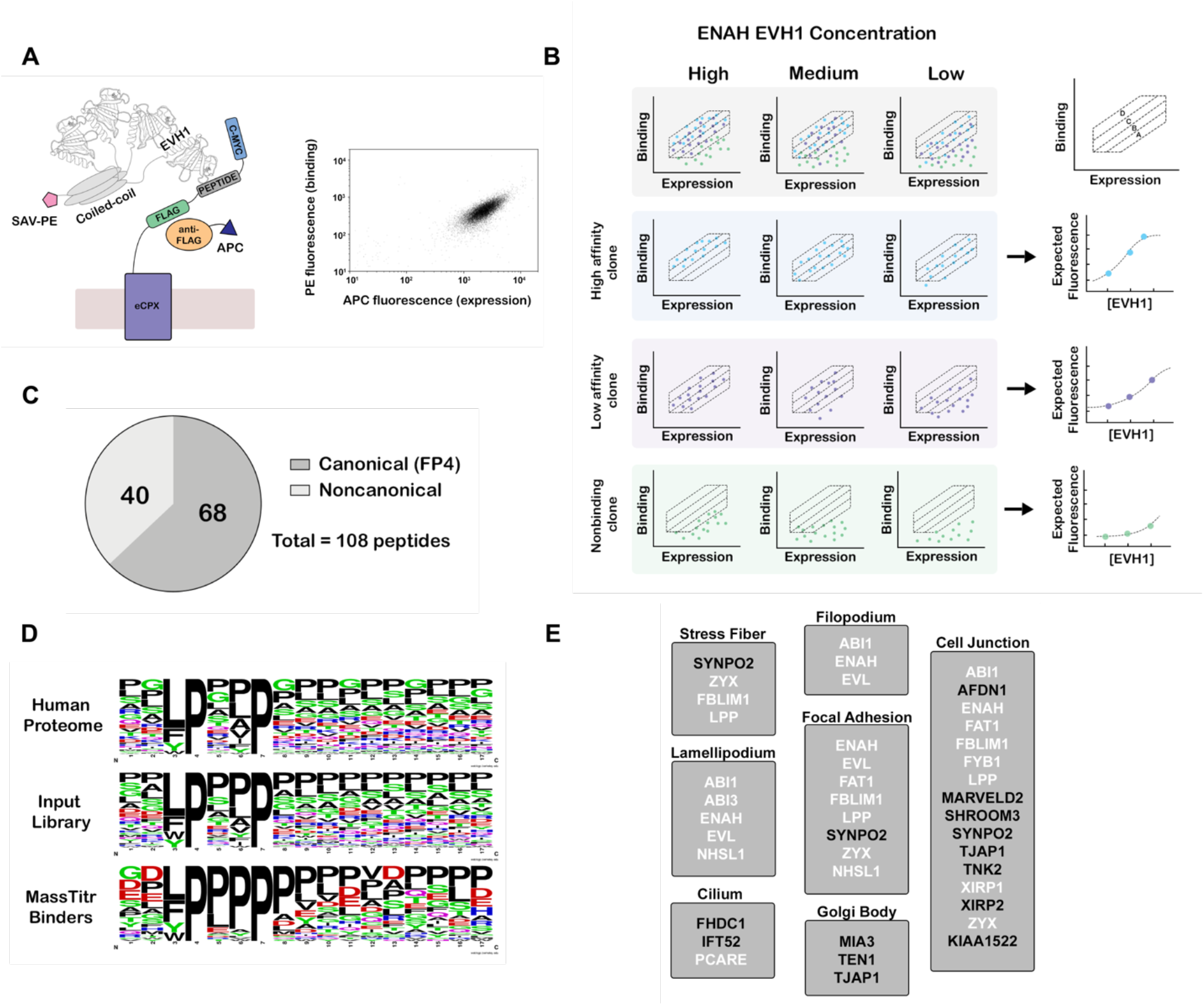
MassTitr screening identifies biologically relevant ENAH EVH1 ligands. (A) At left, bacterial surface display schematic. Library peptides flanked by a FLAG tag and a c-myc tag were expressed as fusions to the C-terminus of eCPX on the surface of *E. coli*. Cells were labeled with anti-FLAG-APC to quantify expression and then incubated with tetrameric ENAH EVH1 domain, which was detected by streptavidin conjugated to phycoerythrin (SAV-PE). At right, a FACS plot for surface-displayed ActA peptide binding to 10 μM ENAH EVH1 tetramer. (B) MassTitr schematic. The top row represents a library of three clones (blue, purple, and green) sorted into four gates at three concentrations of ENAH. The rows highlighted in blue, purple, and green illustrate reconstructions of the concentration-dependent binding of each clone based on deep sequencing. The experiment in this paper sorted a pre-enriched library of clones into four gates at eight concentrations. (C) Distribution of MassTitr hits after filtering; 68 peptides contained a canonical FP4 motif matching the regular expression [FWYL]PX[FWYLIAVP]P. (D) Frequency plot made from the sequences that match the FP4 motif in the human proteome, the input library, and the MassTitr binders using Weblogo (Crooks et al., 2004). (E) Subcellular locations where at least two MassTitr hits that are predicted to be disordered and localized in the cytoplasm are annotated to reside. White text denotes previously reported Ena/VASP interactions.

Using MassTitr, we screened a library of 416,611 36-mer peptides with 7-residue overlaps (the T7-pep library) (Larman et al., 2011). This library spans the entire proteincoding space of the human genome, and we hypothesized that the long lengths of the encoded peptides would illuminate the impact of sequence surrounding the FP4 motif in a biologically relevant sequence space. We first prescreened the library for binding to a tetramerized ENAH EVH1 domain that contained the ENAH EVH1 domain fused to the endogenous ENAH coiled coil as shown in Figure 1A, generating an input library enriched in binders. We then ran MassTitr on the prescreened library, using eight concentrations of ENAH EVH1 tetramer. After sorting, sequencing, and filtering based on read counts, 108 unique high-confidence binders were identified (Figure 1C, Table S1) and classified as either high-affinity or low-affinity as described in the methods.

We validated binding of 16 MassTitr peptide hits to monomeric ENAH EVH1 domain, using biolayer interferometry (BLI) to determine dissociation constants that ranged from 0.19 μM to 63 μM (Table S2). Except for SHROOM3 and TENM1 peptides, binders classified as high-affinity by MassTitr bound to ENAH EVH1 domain more tightly than peptides classified as low-affinity. Many newly identified peptides bound with affinities similar to or tighter than a well-studied control peptide from *Listeria monocytogenes* protein ActA, which bound with K_D_ = 5.2 μM in our BLI assay (Table S2). Prior to this work, this single FP4-motif-containing sequence from ActA was the tightest known endogenously derived binder of Ena/VASP EVH1 domains (Ball et al., 2000). The highest affinity peptide that we discovered was from cilia-associated protein PCARE (K_D_ = 0.19 μM for 36-residue peptide PCARE^813-848^; Table S2), which contains the FP4 motif LPPPP. Successive truncations of this peptide identified the 23-residue minimal region for high-affinity binding, which extends 14 residues beyond the FP4 motif (PCARE^826-848^, which we call PCARE B, K_D_ = 0.32 μM, Figure S1).

Although the majority of MassTitr hits contained FP4 motifs (Figures 1C, D), 40 out of the 108 high-confidence hits did not (Figure 1C). We confirmed that several of these noncanonical peptides (no FP4 motif) bind reversibly to the ENAH EVH1 domain with mid-micromolar affinity (Table S2, Figure S2). Our results support increasing evidence that the ENAH EVH1 domain can bind sequences beyond the FP4 motif (Boëda et al., 2007; Chen et al., 2014; Menon et al., 2015).

### MassTitr peptides are associated with and expand the ENAH signaling network

To highlight putative biologically relevant interaction partners of ENAH, we applied a bioinformatic analysis to identify those motifs that are likely to be accessible and colocalized with ENAH. We filtered our high-confidence hits by disorder propensity (IUPred2A > 0.4) (Mészáros et al., 2018) and cytoplasmic subcellular localization (Binns et al., 2009; Thul et al., 2017). This resulted in 33 peptides, of which 14 are derived from interaction partners previously known to interact or co-localize with an Ena/VASP protein (Table S1). Filtered hits were highly enriched in GO biological process terms including actin filament organization (FDR < 10^-6^), and positive regulation of cytoskeleton (FDR < .05) (Mi et al., 2019), which align with documented cellular functions of ENAH. Notably, we also identified proteins localized to the Golgi body and cilia, where Ena/VASP function is not well characterized (Figure 1E) (Kannan et al., 2014; Tang et al., 2016).

### A proline-rich C-terminal flank binds to a novel site on the EVH1 domain to enhance affinity in ENAH interaction partners

We used MassTitr data to identify FP4 SLiM-flanking elements that enhance binding to the ENAH EVH1 domain. A sequence logo made of the high-confidence MassTitr hits shows enrichment of prolines C-terminal to the FP4 motif, and a binomial test confirms that peptides containing FP4 motifs followed by three consecutive prolines are enriched our hit list (p < 10^-11^; Figure 1D). A peptide from ENAH interactor ABI1 (Chen et al., 2014; Tani et al., 2003) was among the highest affinity ligands that we validated by BLI, with K_D_ = 4 μM (Table S2). ABI1 contains an FP4 motif followed by 4 prolines (referred to here as FP_8_). Mutating FPPPPPPPP to FPPPPSSSS in the context of the ABI1 36-mer reduced affinity by approximately 4-fold (Table S3). Although this confirms that the C-terminal prolines enhance affinity, peptide FPPPPPPPP alone binds to the ENAH EVH1 domain with K_D_ = 29 μM, indicating that additional interactions contribute to the high affinity of the ABI1 36-mer. Previous studies have shown that acidic residues N- and C-terminal to the FP4 motif can enhance affinity (Niebuhr et al., 1997) and we hypothesized that positively charged patches in ENAH could bind acidic residues that flank the FP_8_ segment in ABI1 (Figure S3). Indeed, truncating the N-terminal or C-terminal acidic flanks of the 36-residue ABI1 peptide decreased affinity by 3- or 2.5-fold, respectively (Table S3).

We solved a crystal structure of the ENAH EVH1-ABI1 peptide complex at 1.88 Å resolution. Only the FP_8_ region was fully resolved in the structure under the crystallization conditions, which included high salt and low pH (Figure 2A). The peptide folds into a PPII helix with prolines 1, 4, and 7 (^0^F**P**PP**P**PP**P**P^8^) contacting the EVH1 surface (Figure 2A, B). The FP4 portion of the peptide binds the canonical FP4 groove, as observed in other structures (Prehoda et al., 1999; Fedorov et al., 1999) whereas the 7^th^ proline docks into a previously uncharacterized site on ENAH composed of Ala12, Phe32, and the aliphatic part of the side chain of Asn90 (Figure 2C). Notably, a similar binding site at the analogous location is used by the Homer EVH1 domain to bind the phenylalanine of PPXXF motifs (Beneken et al., 2000). However, this site is relatively shallow in ENAH, and modeling large aromatic acids at this position on the ABI1 peptide using Pymol leads to severe steric clashes. Homer contains a smaller Gly89 at the site of Asn90 in ENAH and can consequently accommodate the bulky Phe of the PPXXF motif (Figure 2C). ENAH paralogs VASP and EVL contain threonine and serine, respectively, in place of Ala12, potentially explaining their ~5-fold weaker affinity for ABI1 (Figure 2C, Table S4). These results reveal how individual EVH1 family members have evolved distinct pockets to recognize proline-rich sequences that extend beyond the core SLiM.

**Figure 2.**
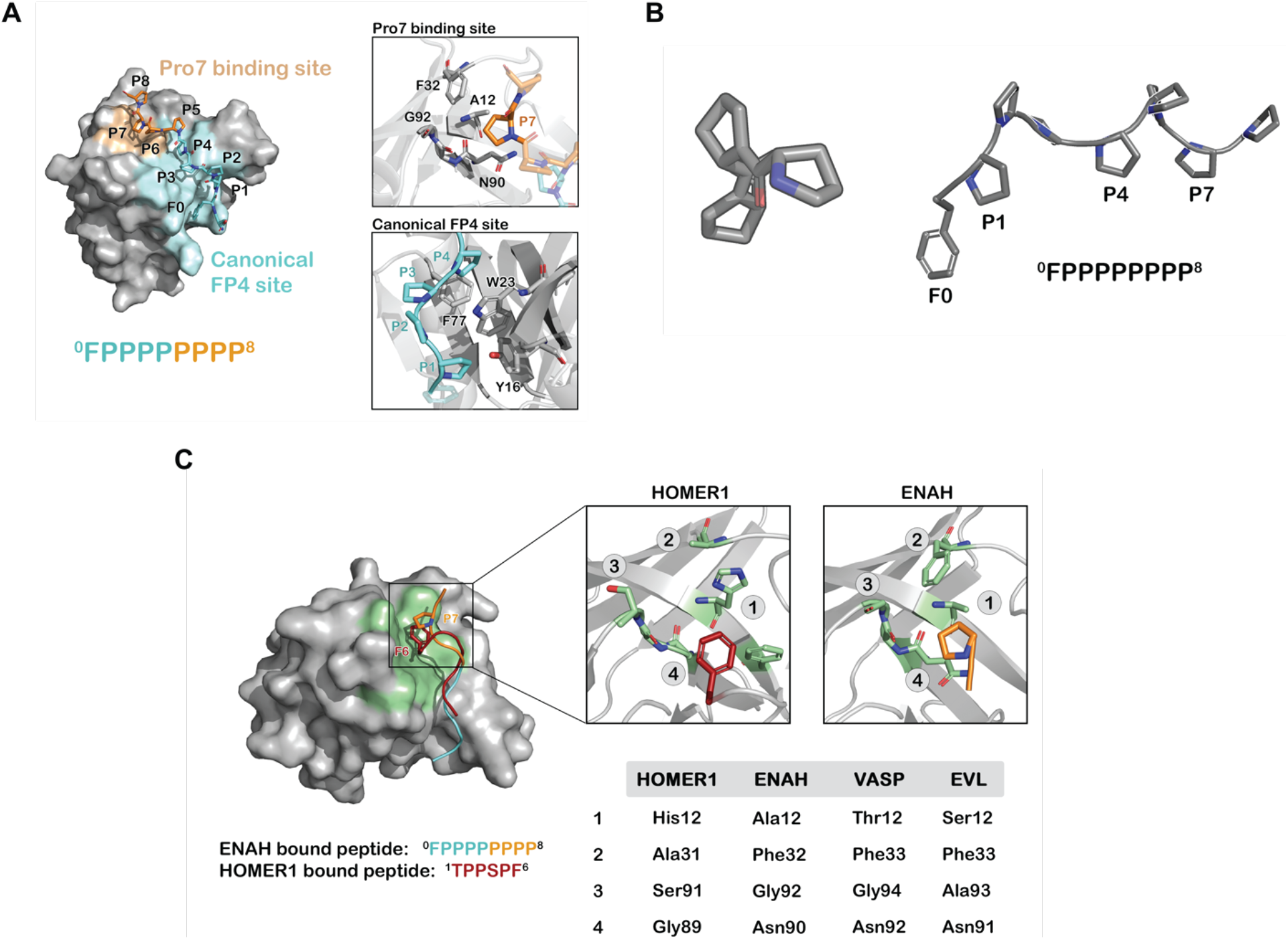
Prolines C-terminal to FP4 can engage a novel ENAH binding site. (A) Surface representation of the ENAH EVH1 domain bound to FP_8_. The core FP4 motif is light blue, the P4 flank is orange; insets show details of the interactions. (B) Axial view of a polyproline type II helix highlighting three-fold symmetry (left); a side view shows P1, P4, and P7 facing the same side (right). (C) At left, surface representation of the HOMER1 EVH1 domain (PDB 1DDV) aligned to the ENAH EVH1 domain, bound to peptide FP_8_ or TPPSPF. The region corresponding to the Pro7 binding pocket in HOMER1 is colored in green. Inset: magnified regions of the Pro7 binding pocket in ENAH and the analogous pocket in HOMER1. The table compares residues in this pocket for HOMER1, ENAH, VASP, and EVL.

### Distal sequence elements enhance ENAH EVH1 binding through bivalent interactions

Another enriched feature of MassTitr-identified binders, relative to the pre-screened input library, is the presence of multiple FP4 motifs (binomial test, p <10^-22^). Multi-motif hits highlight preferred spacings of approximately 5 or 15 residues between FP4 motifs (Figure 3A). Multiple motifs are also enriched in MassTitr high-affinity hits relative to all hits (p < 0.02), supporting the hypothesis that multiple FP4 motifs enhance affinity. We confirmed this experimentally by showing that binding was reduced by 3-6 fold when 36-mer sequences from LPP and NHSL1, which contain two FP4 motifs, were truncated to leave only one motif (Table 1, Figure 3B).

**Figure 3.**
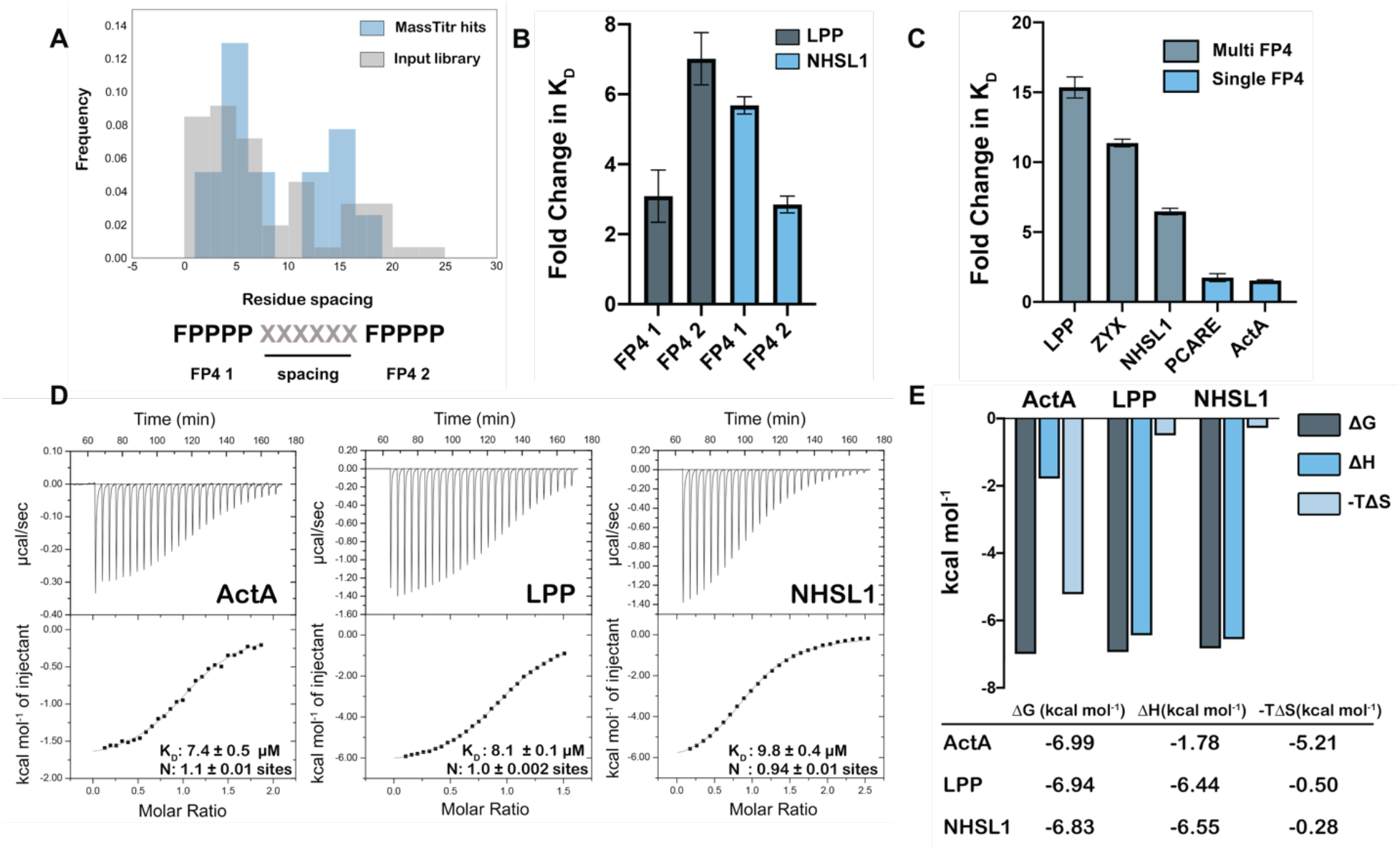
Multiple FP4 motifs enhance peptide binding affinity. (A) Spacing of FP4 motifs in the input library and in high-confidence hits. (B) Fold change in K_D_ for peptide variants relative to tighter-binding 36-mer library peptides for LPP, NHSL1, or zyxin; see Table 1 for sequences. (C) Fold change in K_D_ for 36-mer peptides binding to ENAH EVH1 R47A relative to tighter-binding ENAH EVH1 WT. (D) ITC binding curves for 36-residue peptides from ActA, LPP, and NHSL1; (E) The entropic and enthalpic contributions to binding determined using data in panel D. Fold-change errors in (B) and (C) were calculated by propagating the error from two affinity measurements. Sequences for peptides referenced in this figure are given in Tables 1 and S5.

**Table 1.**
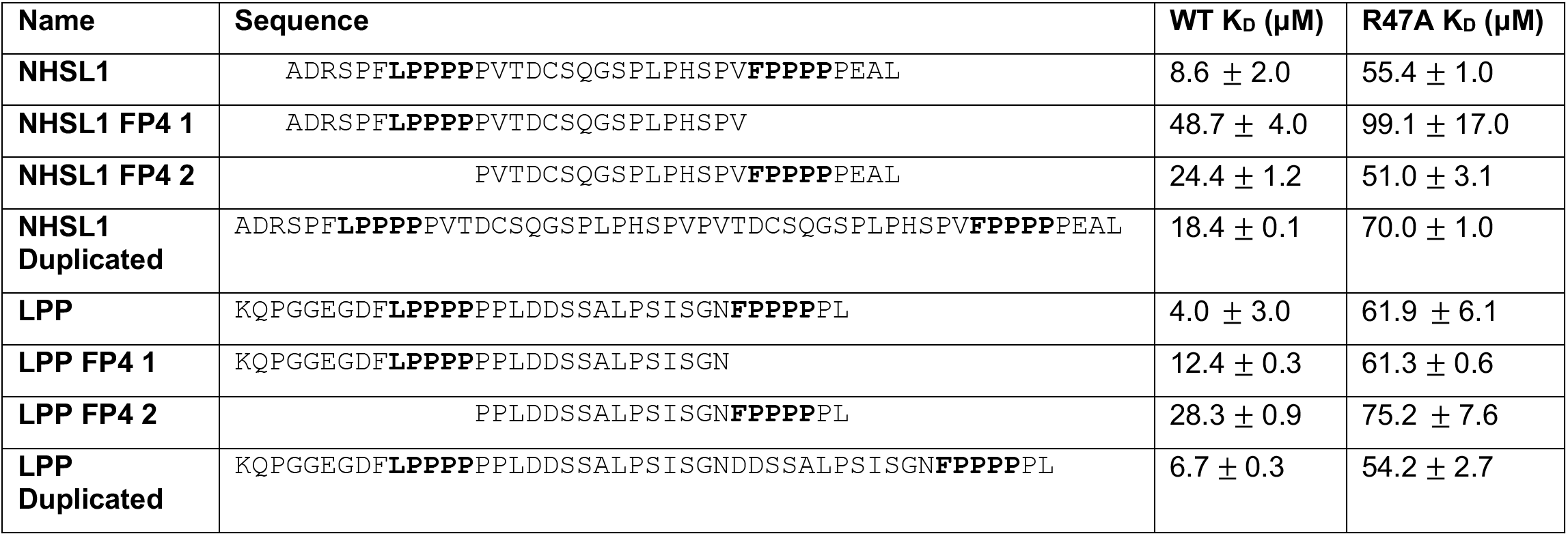
Affinities of dual FP4 motif peptides and their variants for ENAH EVH1 WT or ENAH EVH1 R47A.

Zyxin, which contains four clustered FP4 motifs, has been shown to bind to the VASP EVH1 domain by contacting both the canonical FP4 site and a noncanonical site on the opposite side of the EVH1 domain (Acevedo et al., 2017). Interestingly, a crystal structure of the ENAH EVH1 domain bound to a single-FP4 motif peptide at the canonical site also contains a second single-FP4 peptide bound to the region corresponding to the noncanonical binding site in VASP (PDB 5NC7, Barone, 2020). To test whether multi-FP4 peptides engage this noncanonical site, we designed ENAH EVH1 R47A. In the ActA peptide-bound structure (PDB 5NC7), ENAH Arg47 forms a bidentate hydrogen bond with a carbonyl on the PPII helix backbone of a single FP4-motif peptide. In addition, the analogous VASP Arg48 exhibits significant NMR HSQC chemical shifts upon titration with a multi-FP4 motif zyxin peptide (Acevedo et al., 2017). Thus, we predicted that the R47A mutation would disrupt back-site binding. Indeed, while the affinities of single-FP4 motif peptides from ActA and PCARE were minimally affected by this mutation, peptides containing two FP4 motifs from zyxin, LPP, and NHSL1 experienced a 7-15-fold reduction in affinity to ENAH R47A relative to wild type (Table S5, Figure 3C).

Next, we investigated the stoichiometry of multi-motif peptide binding. Using isothermal titration calorimetry (ITC), we confirmed that dual-FP4 motif peptides from LPP and NHSL1 and the single-FP4 motif peptide from ActA fit well to a 1:1 binding model (Figure 3D). Interestingly, the ITC analysis showed that binding of the ActA-derived single-FP4 motif peptide was driven by favorable entropy, whereas binding of the NHSL1 and LPP dual-motif peptides was enthalpically driven. ActA, LPP, and NHSL1 peptides have similar binding free energies, but the entropic contribution to the dual-motif interactions is ~10-fold less favorable (Figure 3E). These data are consistent with a model in which long, disordered dual-motif peptides pay an entropic penalty to wrap around the EVH1 domain and engage two sites but gain enthalpic binding energy from additional interactions across the EVH1 interface.

Interestingly, duplication of the linker between LPP and NHSL1 led to only a ~2-fold reduction in affinity to the ENAH EVH1 domain compared to the WT LPP and NHSL1 peptide (Table 1). This suggests that linker residues may make compensating interactions with the noncanonical site of the ENAH EVH1 domain.

Finally, we examined the minimal motif-spacing requirements of bivalent binding. We used Rosetta to build a peptide chain to connect single FP4-motif peptides bound to the canonical and noncanonical sites of the ENAH EVH1 domain. There are two orientations that preserve the directionalities of the bound FP4 peptides observed in structure 5NC7 (Figure S4). Ten residues were required to span the two motifs in orientation 1, whereas 9 residues were sufficient in orientation 2 (Figure S4).

## Discussion

In recent years, phage display screening of peptides derived from the human proteome have been used to define SLiM specificity profiles and predict novel interaction partners (Davey et al., 2017; Ueki et al., 2019; Jespersen et al., 2019; Wigington et al., 2020); these studies have primarily focused on defining the “core SLiM”. In this work, we used MassTitr to screen more than 400,000 36-residue segments of the human proteome against the cytoskeleton regulator ENAH and identified both local and distal sequence features up to 15 residues away from the core FP4 SLiM that are important for binding. Our study highlights ways in which low-information SLiMs exploit sequence context to selectively recognize modular interaction domains within the proteome, especially in the context of proline-rich motif-signaling networks where over 300 SH3 domains, 80 WW domains, and 20 EVH1 domains coexist to drive signal transduction in humans (Zarrinpar et al., 2003).

We found multiple ways that sequence flanking the FP4 motif can modulate binding to increase specificity to ENAH. We first demonstrated that prolines C-terminal to FP4 motifs can enhance binding by contacting a previously uncharacterized hydrophobic patch on ENAH. Both secondary structure and sequence are key to this binding mode, which positions the 7^th^ proline of a ^0^FPPPPPP**P**P^8^ peptide to contact ENAH in a shallow groove that we refer to as the Pro7 binding pocket. The relatively flat surface of the EVH1 domain in this region limits the binding energy available from favorable contacts, but PPII helix preorganization minimizes the entropic cost of binding. We anticipate that this binding mode is widely exploited by ENAH cellular interaction partners of ENAH. Multiple previously annotated Ena/VASP interactors, including proteins identified in our screen such as FBLIM1, ZYX, and LPP (Zhang et al., 2006; Drees et al., 2000; Petit et al., 2000) contain FP4 motifs followed by trailing prolines, with either a leucine or proline in the 7^th^ position (^0^FPPPPPP**<u>P</u>**P^8^).

Interestingly, the Homer EVH1 domain uses a site structurally analogous to the ENAH Pro7 binding pocket to accommodate Phe in the SLiM PPXXF (Beneken et al., 2000). Thus, part of the core binding site in the Homer EVH1 domain is used by the ENAH EVH1 domain as a secondary affinity-enhancing site. Ena/VASP family members VASP and EVL bind ~5-fold less tightly than ENAH does to ABI1. These proteins have a polar Thr or Ser in place of an Ala in this pocket, which may account for the specificity. The unique hydrophobic pocket on ENAH EVH1 provides a striking example of how SLiMs can use flanking sequence to target diverged sites in homologous domains to increase molecular recognition specificity.

We also demonstrated that multiple FP4 motifs in ENAH binders can enhance affinity. Many known Ena/VASP partners contain multiple FP4 motifs (Hansen and Mullins, 2015) but the mechanisms by which ENAH engages such peptides are not characterized. Our data and previous work support a model of bivalent binding, where multi-FP4 motif peptides can wrap around a single EVH1 domain (Acevedo et al., 2017). Analysis of our multi-FP4 MassTitr hits showed preferential spacing of ~5 or 15 residues between FP4 motifs in a single chain, but FP4 motifs separated by 5 residues probably do not bind simultaneously to a single EVH1 domain, as structural modeling suggests that the minimum chain length required to span the two putative bindings sites is 9 residues. In such cases, it may be that two EVH1 domains bind to two closely spaced motifs (see one possible model in Figure S2B). Another possibility is that clustered FP4 motifs separated by only a few residues bind using mechanisms such as allovalency, where the increased effective concentration of multiple FP4 motifs close together enhances affinity (Levchenko, 2003).

The sequence requirements for ligands binding to the noncanonical site on ENAH are unknown, but we speculate that diverse sequences, particularly those with the propensity to adopt a PPII helix conformation, could occupy this site when present at high effective concentration due to the binding of a primary motif at the canonical site. In support of this, we saw that peptides from LPP and NHSL1 that lacked a second FP4 motif but contained at least one proline residue ~10 residues away from a single-FP4 motif bound more tightly to wild-type ENAH EVH1 than to ENAH EVH1 R47A (Table 1). Moreover, dual-FP4 peptides from LPP and NHSL1 in which the linker was duplicated bound with similar affinities as the WT peptides to the ENAH EVH1 domain, and also bound less tightly to ENAH EVH1 R47A. This is consistent with a model in which the linker itself can provide stabilizing interactions and the second FP4 motif is not essential for bivalent binding (Table 1). Our data suggest that either a second FP4 motif or linker residues can make favorable interactions with the noncanonical site.

The critical noncanonical site residues for binding FP4 motifs, including ENAH Arg47 and VASP Tyr38 (Acevedo et al., 2017) are conserved across the ENAH, VASP, and EVL, suggesting that bivalent binding is a general mechanism to increase molecular recognition specificity for the Ena/VASP family. However, there is also some evidence that this binding mode could provide paralog specificity, as the linker region connecting multiple FP4 motifs could contact regions on the EVH1 domain that differ across the Ena/VASP paralogs. In support of this, we found that a dual-FP4-motif peptide from LPP bound 8-fold tighter to ENAH over EVL EVH1 domains (Table S4).

Finally, we identified a peptide derived from PCARE that binds to ENAH (K_D_ = 0.19 μM) with the highest known affinity of any SLiM. Truncation experiments of this peptide indicate that the 14-residues C-terminal the LPPPP motif in PCARE are critical for its high affinity, hinting that extensive contacts between this region and the ENAH EVH1 domain could be responsible for the high affinity. Our subsequent work revealed the surprising basis for this affinity (Hwang et al. 2021; co-submission).

Filtering MassTitr hits for interactions of most probable biological significance, based on localization and disorder, yielded peptides from 33 putative binding partners. 19 proteins from this list have, to our knowledge, not been reported to associate with Ena/VASP proteins and provide avenues for further investigation. For example, one of our hits was from the protein KIAA1522, which is known to potentiate metastasis in esophageal carcinoma and breast cancer cells (Xie et al., 2017; Li et al., 2018b). We confirmed that a noncanonical motif in KIAA1522 binds to the ENAH EVH1 domain (K_D_ = 14 μM; Table S2), potentially linking ENAH and KIAA1522 in tumor progression.

Furthermore, several formin proteins (FHOD1, FHDC1, FMN2) were identified as putative ENAH interactors in our screen. Like Ena/VASP proteins, formins also promote unbranched actin polymerization. There is evidence that the two families cooperate in regulating filopodial protrusions (Barzik, 2014), although the mechanistic basis behind this interaction is not well understood. Our hits are potential leads to further investigate the intersection between formins and Ena/VASP proteins in fine-tuning filopodial formation and dynamics. Finally, PCARE, a protein localized to ciliary structures, has been reported to associate with ENAH through tandem affinity purification mass spectrometry and we have now identified the residues that mediate a direct interaction (Corral-Serrano et al., 2020). Investigating the biological implications of the high affinity of PCARE towards ENAH will be of interest.

## Conclusion

For many protein domains beyond EVH1, degenerate SLiMs have been cataloged in the Eukaryotic Linear Motif database to describe their interaction preferences. The ELM listing implies that there is a relatively simple recognition code for many key domain interactions. However, the short sequences of most SLiMs are likely insufficient for biological specificity in many or most cases. Here we showed how defining the EVH1 binding motif as [FWYL]PXΦP is an over-simplification and how, by systematically examining the role of flanking sequences for just one EVH1 domain, we readily uncovered numerous examples in which the affinity and specificity of a core SLiM is modulated via additional extra-motif residues. Added to prior anecdotal examples, our work definitively demonstrates the importance of sequence context on SLiM behavior by illustrating specific mechanisms, including an unusual conformational specificity mechanism that is documented in our companion paper (Hwang et al. 2021 co-submission). MassTitr provides a versatile experimental platform for uncovering context effects on domain-peptide interactions and will surely lead to similar insights into the recognition strategies of other domains.

## Materials and Methods

### FACS sample preparation and analysis

The protocol for sample preparation for FACS analysis and MassTitr sorting was adapted from Foight and Keating (Foight and Keating, 2016) and Whitney et al. (manuscript in preparation) and is as follows: 5 mL cell cultures of eCPX plasmid expressing either library or control peptide were grown overnight at 37 °C in LB + 25 μg/ml chloramphenicol + 0.2% w/v glucose. The next day cells were inoculated into fresh TB + 25 μg/mL chloramphenicol and grown at 37 °C. Upon reaching an O.D. 600 of 0.5-0.6, cells were induced with 0.04% w/v arabinose for 1.5 hours at 37 °C. The O.D. 600 was then remeasured, and enough cells were pelleted for analysis (1 x 10^7^ cells per FACS analysis sample, 7 x 10^7^ cells for library sorting). Cells were resuspended to a concentration of 4 x 10^8^ cells/mL, washed in PBS + 0.1% BSA, and then incubated with anti-FLAG antibody conjugated to APC (αFLAG-APC; PerkinElmer) diluted 1:100 in PBS + 0.1% BSA at a ratio of 30 μL labeled antibody:10^7^ cells. Tubes wrapped in foil were incubated at 4 °C for 15 minutes, then cells were washed with PBS + 0.1% BSA and pelleted. For each FACS analysis sample, 25 μL of 1 x 10^7^ cells in PBS were mixed with 25 μL of a 2x concentration of ENAH tetramer in PBS + 1% BSA + 4 mM DTT (final concentration of 2 mM DTT) and then incubated at 4 °C for 1 hour in foil. After incubation, 50 μL of the mixture was added per well to a 96-well Multi-Screen HTS GV sterile filtration plate (Millipore), buffer was removed by vacuum, and then cells were washed twice with 200 μL of PBS + 0.5% BSA. Each well containing 1 x 10^7^ cells was then resuspended in 30 μL of streptavidin-PE (SAV-PE; ThermoFisher Scientific) diluted 1:100 in PBS + 0.1% BSA, and incubated for 15 minutes at 4 °C, washed with 200 μL of PBS + 0.1% BSA, and resuspended in 250 μL of PBS + 0.1% BSA for subsequent FACS analysis or sorting. For cell sorting see supplementary methods.

### Isothermal titration calorimetry (ITC)

ITC experiments were performed with two replicates using a VP-ITC microcalorimeter (MicroCal LLC). To prepare samples for ITC, 2.5 mL of 100 μM ENAH EVH1 domain and 1 mL of 800 μM-1.2 mM of SUMO-peptide fusions were dialyzed against 2 L of ITC Buffer (20 mM HEPES pH 7.6, 150 mM NaCl, 1 mM TCEP) at 4 °C overnight. The concentrations of proteins were remeasured after dialysis on the day of the experiment. SUMO-peptide was titrated into ENAH EVH1 domain at 25 °C. Data analysis and curve fitting were performed with the Origin 7.0 software (OriginLab). The error reported is the fitting error.

### Crystallography

Crystals of ENAH fused at the C-terminus to ABI1 were grown in hanging drops over a reservoir containing 0.1 M sodium acetate pH 4.5 and 2.90 M NaCl. 1 μL of ENAH-ABI1 (250 μM in 20 mM HEPES, 150 mM NaCl, 1 mM DTT) was mixed with 1 μL of reservoir solution, and 3D crystals appeared in two weeks at 18 °C. Diffraction data were collected at the Advanced Photon Source at Argonne National Laboratory, NE-CAT beamline 24-IDE. The ENAH-ABI1 dataset was integrated and scaled to 1.88 Å with AIMLESS and the structure was solved with molecular replacement using ENAH EVH1 structure 5NC7 as a search model. The structure was refined using iterative rounds of model rebuilding with PHENIX and COOT. Table S7 reports refinement statistics. The structure is deposited in the PDB with identifier 7LXE. Note that the ABI1 FP_8_ peptide is numbered 120-129 in accordance with the ENAH-ABI1 fusion protein numbering.

## Acknowledgments

This project was supported by NIGMS award R01 GM129007 to A.E.K. Part of this work is based upon research conducted at the Northeastern Collaborative Access Team beamlines, which are funded by the National Institute of General Medical Sciences from the National Institutes of Health (P30 GM124165). The Eiger 16M detector on the 24-IDE beamline is funded by an NIH-ORIP HEI grant (S10OD021527). This research used resources of the Advanced Photon Source, a U.S. Department of Energy (DOE) Office of Science User Facility operated for the DOE Office of Science by Argonne National Laboratory under contract DE-AC02-06CH11357. T.H. was partially supported by NIGMS T32 GM007287 and a fellowship from the Koch Institute for Integrative Cancer Research.

We thank the Koch Institute Flow Cytometry Core Facility for assistance with fluorescence-activated cell sorting (FACS), the MIT Structural Biology Core for assistance with X-ray crystallography, the MIT Biophysical Instrumentation Facility for instrumentation resources, and the MIT BioMicroCenter for assistance with high-throughput sequencing. We thank D. Whitney for contributions to the MassTitr technology, and we thank members of the Keating lab and F. Gertler for their thoughtful input. We also thank J. Tadros and F. Gertler for reagents. We also thank M. Li, S. Elledge, and Brigham and Women’s hospital for the T7-pep library.

## Author Contributions

T.H. and A.E.K. designed the study. T.H and M.W.I. performed the experiments and structure modeling. R.A.G. helped with X-ray crystallography experiments. T.H., R.A.G., and M.W.I. analyzed data. V.S. built the pipeline to process next-generation sequencing data. T.H. wrote the original draft. A.E.K. reviewed and edited the paper. A.E.K. provided supervision and funding acquisition. A.E.K. oversaw all experimental and computational work reported here.

## References

1. Adams RM, Mora T, Walczak AM, Kinney JB. Measuring the sequence-affinity landscape of antibodies with massively parallel titration curves. eLife 5, e23156 (2016).

2. Acevedo LA, Greenwood AI, Nicholson LK. A Noncanonical Binding Site in the EVH1 Domain of Vasodilator-Stimulated Phosphoprotein Regulates Its Interactions with the Proline Rich Region of Zyxin. Biochemistry 56, 4626–4636 (2017).

3. Ball, LJ, Kühne R., Hoffmann B, Häfner A, Schmieder P, Volkmer-Engert R, Hof M, Wahl M, Schneider-Mergener J, Walter U, Oschkinat H, Jarchau T. Dual epitope recognition by the VASP EVH1 domain modulates polyproline ligand specificity and binding affinity. EMBO J. 19, 4903–4914 (2000).

4. Barone M, Müller M, Chiha S, Ren J, Albat D, Soicke A, Dohmen S, Klein M, Bruns J, van Dinther M, Opitz R, Lindemann P, Beerbaum M, Motzny K, Roske Y, Schmieder P, Volkmer R, Nazaré M, Heinemann U, Oschkinat H, … Kühne R. Designed nanomolar small-molecule inhibitors of Ena/VASP EVH1 interaction impair invasion and extravasation of breast cancer cells. Proc. Natl. Acad. Sci. U.S.A. 117, 29684–29690 (2020).

5. Barzik M, McClain LM, Gupton SL, Gertler FB. Ena/VASP regulates mDia2-initiated filopodial length, dynamics, and function. Mol. Biol. Cell. 5, 2604–2619 (2014).

6. Beneken J, Tu JC, Xiao B, Nuriya M, Yuan JP, Worley PF, Leahy DJ. Structure of the Homer EVH1 domain-peptide complex reveals a new twist in polyproline recognition. Neuron 26, 143–54 (2000).

7. Bhattacharyya RP, Reményi A, Yeh BJ, Lim WA. Domains, motifs, and scaffolds: the role of modular interactions in the evolution and wiring of cell signaling circuits. Annu. Rev. Biochem. 75, 655–680 (2006).

8. Binns D, Dimmer E, Huntley R, Barrell D, O’Donovan C, Apweiler R. QuickGO: a web-based tool for Gene Ontology searching. Bioinformatics 25, 3045–3046 (2009).

9. Boëda B, Briggs DC, Higgins T, Garvalov BK, Fadden AJ, McDonald NQ, Way M. Tes, a specific Mena interacting partner, breaks the rules for EVH1 binding. Mol. Cell 28, 1071–1082 (2007).

10. Bugge K, Brakti I, Fernandes CB, Dreier JE, Lundsgaard JE, Olsen JG, Skriver K, Kragelund BB. Interactions by Disorder - A Matter of Context. Front. Mol. Biosci. 7, 110 (2020).

11. Chen XJ, Squarr AJ, Stephan R, Chen B, Higgins TE, Barry DJ, Martin MC, Rosen MK, Bogdan S, Way M. Ena/VASP proteins cooperate with the WAVE complex to regulate the actin cytoskeleton. Ena/VASP proteins cooperate with the WAVE complex to regulate the actin cytoskeleton. Dev. Cell 30, 569–584 (2014).

12. Corral-Serrano JC, Lamers IJC, van Reeuwijk J, Duijkers L, Hoogendoorn ADM, Yildirim A, Argyrou N, Ruigrok RAA, Letteboer SJF, Butcher R, van Essen MD, Sakami S, van Beersum SEC, Palczewski K, Cheetham ME, Liu Q, Boldt K, Wolfrum U, Ueffing M, Garanto A, Roepman R, Collin RWJ. PCARE and WASF3 regulate ciliary F-actin assembly that is required for the initiation of photoreceptor outer segment disk formation. Proc. Natl. Acad. Sci. U.S.A. 117, 9922–9931 (2020).

13. Crooks GE, Hon G, Chandonia JM, Brenner SE. WebLogo: a sequence logo generator. Genome Res. 14, 1188–1190 (2004).

14. Davey NE, Seo MH, Yadav VK, Jeon J, Nim S, Krystkowiak I, Blikstad C, Dong D, Markova N, Kim PM, Ivarsson Y. Discovery of short linear motif-mediated interactions through phage display of intrinsically disordered regions of the human proteome. FEBS J. 284, 485–498 (2017).

15. Drees B, Friederich E, Fradelizi J, Louvard D, Beckerle MC, Golsteyn RM. Characterization of the interaction between zyxin, and members of the Ena/vasodilator-stimulated phosphoprotein family of proteins. J. Biol. Chem. 275, 22503–22511 (2000).

16. Fedorov AA, Fedorov E, Gertler F, Almo SC. Structure of EVH1, a novel proline-rich ligand-binding module involved in cytoskeletal dynamics and neural function. Nat. Struct. Mol. Biol. 6, 661–665 (1999).

17. Foight GW, Keating AE. Comparison of the peptide binding preferences of three closely related TRAF paralogs: TRAF2, TRAF3, and TRAF5. Protein Sci. 25, 1273–1289 (2016).

18. Hansen SD, Mullins RD. Lamellipodin promotes actin assembly by clustering Ena/VASP proteins and tethering them to actin filaments. eLife 4, e06585 (2015).

19. Holt MR, Koffer A. Cell motility: proline-rich proteins promote protrusions. Trends Cell Biol. 11, 38–46 (2001).

20. Ivarsson Y, Arnold R, McLaughlin M, Nim S, Joshi R, Ray D, Liu B, Teyra J, Pawson T, Moffat J, Li SS, Sidhu SS, Kim PM. Large-scale interaction profiling of PDZ domains through proteomic peptide-phage display using human and viral phage peptidomes. Proc. Natl. Acad. Sci. U.S.A. 111, 2542–2547 (2014).

21. Jespersen N, Estelle A, Waugh N, Davey NE, Blikstad C, Ammon YC, Akhmanova A, Ivarsson Y, Hendrix DA, Barbar E. Systematic identification of recognition motifs for the hub protein LC8. Life Sci. Alliance 2, e201900366 (2019).

22. Kannan R, Kuzina I, Wincovitch S, Nowotarski SH, Giniger E. The Abl/enabled signaling pathway regulates Golgi architecture in Drosophila photoreceptor neurons. Mol. Biol. Cell 25, 2993–3005 (2014).

23. Larman HB, Zhao Z, Laserson U, Li MZ, Ciccia A, Gakidis MA, Church GM, Kesari S, Leproust EM, Solimini NL, Elledge SJ. Autoantigen discovery with a synthetic human peptidome. Nat. Biotechnol. 29, 535–541 (2011).

24. Levchenko A. Allovalency: a case of molecular entanglement. Curr. Biol. 13, R876–R878 (2003).

25. Li J, Zhu R, Chen K, Zheng H, Zhao H, Yuan C, Zhang H, Wang C, Zhang M. Potent and specific Atg8-targeting autophagy inhibitory peptides from giant ankyrins. Nat Chem Biol. 14, 778–787 (2018a).

26. Li Y, Wang Y, Fan H, Zhang Z, Li N. miR-125b-5p inhibits breast cancer cell proliferation, migration and invasion by targeting KIAA1522. Biochem. Biophys. Res. Commun. 504, 277–282 (2018b).

27. Li Z, Liu H, Li J, Yang Q, Feng Z, Li Y, Yang H, Yu C, Wan J, Liu W, Zhang M. Homer Tetramer Promotes Actin Bundling Activity of Drebrin. Structure 27, 27–38 (2019).

28. Lim WA, Richards FM, Fox RO. Structural determinants of peptide-binding orientation and of sequence specificity in SH3 domains. Nature 372, 375–379 (1994).

29. Macias MJ, Hyvönen M, Baraldi E, Schultz J, Sudol M, Saraste M, Oschkinat H. Structure of the WW domain of a kinase-associated protein complexed with a proline-rich peptide. Nature 382, 646–649 (1996).

30. McConnell RE, Edward van Veen J, Vidaki M, Kwiatkowski AV, Meyer AS, Gertler FB. A requirement for filopodia extension toward Slit during Robo-mediated axon repulsion. J. Cell Biol. 213, 261–274 (2016).

31. Menon S, Boyer NP, Winkle CC, McClain LM, Hanlin CC, Pandey D, Rothenfußer S, Taylor AM, Gupton SL. The E3 Ubiquitin Ligase TRIM9 Is a Filopodia Off Switch Required for Netrin-Dependent Axon Guidance. Dev. Cell 35, 698–712 (2015).

32. Mészáros B, Erdos G, Dosztányi Z. IUPred2A: context-dependent prediction of protein disorder as a function of redox state and protein binding. Nucleic Acids Res. 46, W329–W337 (2018).

33. Mi H, Muruganujan A, Huang X, Ebert D, Mills C, Guo X, Thomas PD. Protocol Update for large-scale genome and gene function analysis with the PANTHER classification system (v.14.0). Nat. Protoc. 14, 703–721 (2019).

34. Niebuhr K, Ebel F, Frank R, Reinhard M, Domann E, Carl UD, Walter U, Gertler FB, Wehland J, Chakraborty T. A novel proline-rich motif present in ActA of Listeria monocytogenes and cytoskeletal proteins is the ligand for the EVH1 domain, a protein module present in the Ena/VASP family. EMBO J. 16, 5433–5444 (1997).

35. Palopoli N, González Foutel NS, Gibson TJ, Chemes LB. Short linear motif core and flanking regions modulate retinoblastoma protein binding affinity and specificity. Protein Eng. Des. Sel. 31, 69–77 (2018).

36. Petit MM, Fradelizi J, Golsteyn RM, Ayoubi TA, Menichi B, Louvard D, Van de Ven WJ, Friederich E. LPP, an actin cytoskeleton protein related to zyxin, harbors a nuclear export signal and transcriptional activation capacity. Mol. Biol. Cell 11, 117–129 (2000).

37. Prehoda KE, Lee DJ, Lim WA. Structure of the enabled/VASP homology 1 domain-peptide complex: a key component in the spatial control of actin assembly. Cell 97, 471–480 (1999).

38. Prestel A, Wichmann N, Martins JM, Marabini R, Kassem N, Broendum SS, Otterlei M, Nielsen O, Willemoës M, Ploug M, Boomsma W, Kragelund BB. The PCNA interaction motifs revisited: thinking outside the PIP-box. Cell. Mol. Life Sci. 76, 4923–4943 (2019).

39. Schultz J, Hoffmüller U, Krause G, Ashurst J, Macias MJ, Schmieder P, Schneider-Mergener J, Oschkinat H. Specific interactions between the syntrophin PDZ domain and voltage-gated sodium channels. Nat. Struct. Mol. Biol. 5, 19–24 (1998).

40. Scott JA, Shewan AM, den Elzen NR, Loureiro JJ, Gertler FB, Yap AS. Ena/VASP proteins can regulate distinct modes of actin organization at cadherin-adhesive contacts. Mol. Biol. Cell 17, 1085–1095 (2006).

41. Tang D, Zhang X, Huang S, Yuan H, Li J, Wang Y. Mena-GRASP65 interaction couples actin polymerization to Golgi ribbon linking. Mol. Biol. Cell 27, 137–152 (2016).

42. Tani K, Sato S, Sukezane T, Kojima H, Hirose H, Hanafusa H, Shishido T. Abl interactor 1 promotes tyrosine 296 phosphorylation of mammalian enabled (Mena) by c-Abl kinase. J. Biol. Chem. 278, 21685–21692 (2003).

43. Teyra J, Huang H, Jain S, Guan X, Dong A, Liu Y, Tempel W, Min J, Tong Y, Kim PM, Bader GD, Sidhu SS. Comprehensive Analysis of the Human SH3 Domain Family Reveals a Wide Variety of Non-canonical Specificities. Structure 25, 1598–1610 (2017).

44. Thul PJ, Åkesson L, Wiking M, Mahdessian D, Geladaki A, Ait Blal H, Alm T, Asplund A, Björk L, Breckels LM, Bäckström A, Danielsson F, Fagerberg L, Fall J, Gatto L, Gnann C, Hober S, Hjelmare M, Johansson F, Lee S, Lindskog C, Mulder J, Mulvey CM, Nilsson P, Oksvold P, Rockberg J, Schutten R, Schwenk JM, Sivertsson Å, Sjöstedt E, Skogs M, Stadler C, Sullivan DP, Tegel H, Winsnes C, Zhang C, Zwahlen M, Mardinoglu A, Pontén F, von Feilitzen K, Lilley KS, Uhlén M, Lundberg E. A subcellular map of the human proteome. Science 356, eaal3321 (2017).

45. Tonikian R, Zhang Y, Sazinsky SL, Currell B, Yeh JH, Reva B, Held HA, Appleton BA, Evangelista M, Wu Y, Xin X, Chan AC, Seshagiri S, Lasky LA, Sander C, Boone C, Bader GD, Sidhu SS. A specificity map for the PDZ domain family. PLOS Biol. 6, e239 (2008).

46. Ueki Y, Kruse T, Weisser MB, Sundell GN, Larsen MSY, Mendez BL, Jenkins NP, Garvanska DH, Cressey L, Zhang G, Davey N, Montoya G, Ivarsson Y, Kettenbach AN, Nilsson J. A Consensus Binding Motif for the PP4 Protein Phosphatase. Mol. Cell 76, 953–964 (2019).

47. Wigington CP, Roy J, Damle NP, Yadav VK, Blikstad C, Resch E, Wong CJ, Mackay DR, Wang JT, Krystkowiak I, Bradburn DA, Tsekitsidou E, Hong SH, Kaderali MA, Xu SL, Stearns T, Gingras AC, Ullman KS, Ivarsson Y, Davey NE, Cyert MS. Systematic Discovery of Short Linear Motifs Decodes Calcineurin Phosphatase Signaling. Mol. Cell 79, 342–358 (2020).

48. Xie ZH, Yu J, Shang L, Zhu YQ, Hao JJ, Cai Y, Xu X, Zhang Y, Wang MR. KIAA1522 overexpression promotes tumorigenicity and metastasis of esophageal cancer cells through potentiating the ERK activity. OncoTargets Ther. 10, 3743–3754 (2017).

49. Zarrinpar A, Bhattacharyya RP, Lim WA. The structure and function of proline recognition domains. Sci. Signal 2003, RE8 (2003).

50. Zhang Y, Tu Y, Gkretsi V, Wu C. Migfilin interacts with vasodilator-stimulated phosphoprotein (VASP) and regulates VASP localization to cell-matrix adhesions and migration. J. Biol. Chem. 281, 12397–12407 (2006).

