## Supplementary Materials for "Native proline-rich motifs exploit sequence context to target actin-remodeling Ena/VASP proteins"

### Table of contents:

1. Protein Constructs
2. Supplementary Materials and Methods
3. Figure S1: **PCARE truncations**
4. Figure S2: **Peptides containing CXC motifs bind to ENAH**
5. Figure S3: **Additional analyses of the ENAH EVH1-ABI1 structure**
6. Figure S4: **Modeling bivalent binding**
7. Figure S5: **Gates drawn for MassTitr FACS Sort**
8. Table S2: **Dissociation constants of MassTitr peptides derived from human proteins to ENAH**
9. Table S3: **ABI1 truncation experiments**
10. Table S4: **Dissociation constants of select peptides to VASP and EVL**
11. Table S5**: Comparison of affinities between ENAH WT or ENAH R47A against multi or single FP4 motif peptides.**
12. Table S7: **Refinement table for ENAH-ABI1 structure**

### Other supplementary materials:

Table S1: **MassTitr hits and annotated interactors.** Provided as an excel spreadsheet. The MassTitr hits tab lists all high-confidence hits from the screen as well as the human protein the peptide sequence corresponds to. The putative interactors tab is a list of MassTitr hits filtered for disorder propensity and subcellular localization that represent putative ENAH interactors.
Table S6: **MassTitr sample preparation and key**. Provided as an excel spreadsheet. The illumina amplicon preparation tab displays the schematic used to prepare amplicons for deep sequencing after MassTitr sorting. The barcode index key tab lists the 5-nucleotide barcode and 6-nucleotide index used to label each sample for multiplexing. The cell counts tab lists the number of cells collected for each MassTitr replicate.

### Protein Constructs

**pDW363 constructs**

Constructs used for bacterial display

**ENAH tetramer**

MSEQSICQARAAVMVYDDANKKWVPAGGSTGFSRVHIYHHTGNNTFRVVGRKIQDH
QVVINCAIPKGLKYNQATQTFHQWRDARQVYGLNFGSKEDANVFASAMMHALEVLNS
QEAGGGGGGGSACEGLDYDRLKQDILDEMRKELAKLKEELIDAIRQELSKSNTAGSGS
GSGLNDIFEAQKIEWHEDTGGSSHHHHHHG*

**Cysteine-less ENAH tetramer**

MSEQSIAQARAAVMVYDDANKKWVPAGGSTGFSRVHIYHHTGNNTFRVVGRKIQDHQ
VVINSAIPKGLKYNQATQTFHQWRDARQVYGLNFGSKEDANVFASAMMHALEVLNSQ
EAGGGGGGGSASEGLDYDRLKQDILDEMRKELAKLKEELIDAIRQELSKSNTAGSGSG
SGLNDIFEAQKIEWHEDTGGSSHHHHHHG*

SUMO-peptide fusions for ITC and BLI

**SUMO only (empty)**

MAGGLNDIFEAQKIEWHEDTGGSSHHHHHHGSGSGSDSEVNQEAKPEVKPEVKPET
HINLKVSDGSSEIFFKIKKTTPLRRLMEAFAKRQGKEMDSLTFLYDGIEIQADQTPEDLD
MEDNDIIEAHREQIGG*

**SUMO-ActA**

MAGGLNDIFEAQKIEWHEDTGGSSHHHHHHGSGSGSDSEVNQEAKPEVKPEVKPET
HINLKVSDGSSEIFFKIKKTTPLRRLMEAFAKRQGKEMDSLTFLYDGIEIQADQTPEDLD
MEDNDIIEAHREQIGGGFNAPATSEPSSFEFPPPPTEDELEIIRETASSLDS*

**All other peptides used in the experiment were cloned into the C-terminus of the following sequence:**

MAGGLNDIFEAQKIEWHEDTGGSSHHHHHHGSGSGSDSEVNQEAKPEVKPEVKPET
HINLKVSDGSSEIFFKIKKTTPLRRLMEAFAKRQGKEMDSLTFLYDGIEIQADQTPEDLD
MEDNDIIEAHREQIGGSGSG[PEPTIDE SEQUENCE]

Peptide sequences inserted after the C-terminus are listed in Tables 1, S2, and S3

**pMCSG7 constructs**

Constructs used for BLI

**ENAH EVH1**

MHHHHHHSSGVDLGTENLYFQSNAMSEQSICQARAAVMVYDDANKKWVPAGGSTGF
SRVHIYHHTGNNTFRVVGRKIQDHQVVINCAIPKGLKYNQATQTFHQWRDARQVYGLN
FGSKEDANVFASAMMHALEVL*

**VASP EVH1**

MHHHHHHSSGVDLGTENLYFQSNAMSETVICSSRATVMLYDDGNKRWLPAGTGPQA
FSRVQIYHNPTANSFRVVGRKMQPDQQVVINCAIVRGVKYNQATPNFHQWRDARQV
WGLNFGSKEDAAQFAAGMASALEALE*

**EVL EVH1**

MHHHHHHSSGVDLGTENLYFQSNAMSEQSICQARASVMVYDDTSKKWVPIKPGQQG
FSRINIYHNTASSTFRVVGVKLQDQQVVINYSIVKGLKYNQATPTFHQWRDARQVYGL
NFASKEEATTFSNAMLFALNIMNSQE*

Constructs used for crystallography

**ENAH EVH1-ABI1**

MHHHHHHSSGVDLGTENLYFQSNAMSEQSICQARAAVMVYDDANKKWVPAGGSTGF
SRVHIYHHTGNNTFRVVGRKIQDHQVVINCAIPKGLKYNQATQTFHQWRDARQVYGLN
FGSKEDANVFASAMMHALEVLGGSGSGFDDFPPPPPPPPVDYEDEEAAVVQYNDPY
ADGDPAW*

**eCPX constructs**

**All peptide sequences, and the T7-pep library, were inserted into the following background:**

MKKIACLSALAAVLAFTAGTSVAGGQSGQSGDYNKNQYYGITAGPAYRINDWASIYGV
VGVGYGKFQTTEYPTYKHDTSDYGFSYGAGLQFNPMENVALDFSYEQSRIRSVDVGT
WILSVGYRFGSKSRRATSTVTGGYAQSDAQGQMNKMGGFNLKYRYEEDNSPLGVIG
SFTYTEKSRTASGGGSGGGSDYKDDDDKGGGSGGGSG**IPLR**[PEPTIDE
SEQUENCE]**RIAR**GSGSEQKLISEEDL*

Bold sequences IPLR and RIAR constitute part of the EcoRI and XhoI restriction enzyme sites, respectively.

Peptide sequences are as follows:

| Name | Sequence |
|---|---|
| Empty | No peptide inserted |
| ActA | GFNAPATSEPSSFEFPPPPTEDELEIIRETASSLDS |
| Vinculin | EAFQPQEPDFPPPPPDLEQLRLTDELAPPKPPLPEG |
| SHIP2 | VGEGSSSDEESGGTLPPPDFPPPPLPDSAIFLPPSL |
| OLIG3 | HWAGLPCPCTICQMPPPPHLSALSTANMARLSAESK |
| TRIM1 | ILSGLPAPDFIDYPERQECNCRPQESPYVSGMKTCH* |

### Supplementary Materials and Methods

Biolayer interferometry, monomeric EVH1 domain purification, and small-scale SUMO-peptide purification as described in Hwang et al., 2021.

**Protein expression and purification**

Human ENAH EVH1 domain, followed by a 6x-Gly linker and ENAH mouse coiled coil (for tetramerization), were cloned into a pDW363 biotinylation vector that includes a C-terminal biotin acceptor peptide (BAP) tag and a 6x-His tag. This ENAH tetramer construct was expressed in Rosetta2(DE3) (Novagen) cells in Terrific Broth (TB) with 100 µg/mL ampicillin, 25 µg/mL chloramphenicol, and 0.05 mM D-(+)-biotin (for *in vivo* biotinylation). Cells were grown at 37 °C with shaking to an optical density at 600 nm (O.D. 600) of 0.5-0.7 and then induced with 1 mM IPTG and grown at 37 °C for 5 hours. 1 L of cells were then spun down and resuspended in 25 mL of binding buffer (20 mM Tris pH 8.0, 500 mM NaCl, 5 mM imidazole), and frozen at -80 °C overnight. The next day, pellets were thawed and supplemented with 0.2 mM phenylmethylsulfonyl fluoride (PMSF) protease inhibitor. Cells were sonicated ten times for 30 s followed by 30 s of rest on ice and then centrifuged. The clarified lysate was filtered through a 0.2 µm filter and applied to 2 mL of Ni-nitrilotriacetic (Ni-NTA) acid agarose resin (GoldBio) equilibrated in wash buffer (20 mM Tris pH 8.0, 500 mM NaCl, 20 mM imidazole). The resin was then washed 3 times with 8 mL wash buffer and eluted with 10 mL of elution buffer (20 mM Tris pH 8.0, 500 mM NaCl, 300 mM imidazole). The elution was run through a S75 26/60 size exclusion column equilibrated in gel filtration buffer (20 mM Tris pH 8.0, 150 mM NaCl, 1 mM DTT, 10% glycerol). Purity was verified by SDS-PAGE, and the fractions were pooled, concentrated, and flash frozen at -80 °C.

SUMO-peptide fusions were cloned into a pDW363 vector that appends a BAP sequence and 6x-His tag to the N-terminus of the protein and transformed into Rosetta2(DE3) cells. For ITC experiments, these cells were expressed in TB supplemented with 100 µg/mL ampicillin, grown to an O.D. 600 of 0.5-0.7, and induced with 1 mM IPTG. Induced cultures were purified as described above for the pMCGS7 constructs, with the exception of the TEV cleavage step. Instead, after elution with elution buffer, the sample was directly applied to the S75 26/60 column equilibrated in gel filtration buffer. Fractions were pooled, concentrated, and flash frozen at -80 °C.

**Bacterial cell surface display plasmids and T7-Pep library cloning**

Control peptides for display were expressed at the C-terminus of eCPX in a vector designed by the Daughtery group (Rice and Daugherty, 2008). This construct was modified to include a FLAG tag at the N-terminus of the peptide and a C-myc tag at the C-terminus. The T7-pep library plasmids (gift from Elledge lab, Harvard), were transformed into Pir1 cells (Thermofisher), grown, and miniprepped (Qiagen) to isolate the library plasmid. Plasmids were then cut with EcoRI (NEB)and XhoI (NEB), and the inserts were gel purified, combined, and concentrated with a Zymo Clean and Concentrate column, and eluted with 50 µL of sterile MilliQ Water. To clone the T7-pep library into the eCPX vector, we first grew up 200 mL of an empty eCPX vector in DH5a cells at 37 °C overnight. This culture was miniprepped (Qiagen) and then digested with

EcoI and XhoI at a ratio of 10 units of enzyme:1 μg of vector at 37 °C for 2 hours. The resulting digest was PCR purified and eluted with 40 μL water. This cut vector was then dephosphorylated with Antarctic phosphatase (NEB) at a ratio of 1 μL/1 μg of DNA at 37 °C for 2 hours, followed by 10 min at 65 °C for enzyme inactivation. T7-pep library insert was ligated into the cut eCPX vector using T4 ligase (NEB) at 14 °C overnight. Ligase was subsequently deactivated for 10 minutes at 70 °C, and the ligation reaction was concentrated with Zymo Clean and Concentrate columns (Zymo Research). Each column was eluted with 12.5 μL elution buffer (from kit). The resulting elutions were desalted on a 0.025 μm filter (Millipore) for 15-20 minutes and pooled on ice. Electrocompetent MC1061 cells and 10-20 μL DNA were then mixed and transferred to a cold 2 mm cuvette (BioRad). Each cuvette was pulsed at 2.5 kV, 50 μF, 100 ohms on an electroporator (BioRad), immediately rinsed out with 3 x 1 mL of warm SOC and transferred to a culture tube containing 7 mL warm SOC. Cells were incubated at 37 °C for one hour and then combined. Serial dilutions of the library were plated on LB/chloramphenicol plates to assess transformation efficiency, and the leftover cells were added to 500 mL of LB + 25 μg/mL chloramphenicol + 0.2% w/v sterile-filtered glucose. The library was grown at 37 °C until it reached an O.D. 600 of 2.0 and then frozen as glycerol stocks to use for FACS analysis and sorting.

**Pre-enrichment of T7-Pep library**

The T7-pep library was prepared as described above. 7 x $10^7$ cells expressing the T7-pep library were incubated with a final concentration of 20 μM ENAH tetramer. Cells were sorted on a BDFACS Aria machine. Prior to sorting, a positive (ActA) control and a negative (empty) control were analyzed. A gate to collect cells expressing peptide binders was set that included 0.3% of the negative control and all cells with a greater binding signal, allowing us to enrich moderate-affinity binders. This process was repeated three times on the same day to collect a total of 300,000 cells. These cells were added to warm SOC + 25 μg/mL chloramphenicol and grown at 37 °C overnight. The next day, cells were frozen at -80 °C as glycerol stocks for MassTitr sorting.

**MassTitr sorting scheme**

The pre-enriched T7-Pep Library was induced and prepared as previously described. 7 x $10^7$ cells were incubated with increasing concentrations of ENAH EVH1 tetramer (monomeric concentration: 30, 12, 4.8, 1.9, .77, 0.31, 0.12, 0.049 μM) and then sorted on the BDFACS Aria. Cells were collected in 4 gates that were drawn with boundaries roughly parallel to the binding vs. expression signal slope for a series of positive controls (ActA, SHIP2, Vinculin) (Figure S5). A no-peptide negative control (empty) was run to assess the degree of nonspecific binding at the time of sorting. At each concentration of ENAH EVH1 domain, cells were sorted for approximately 20 minutes and the number of cells that were collected in each gate was recorded. Enough cells were collected per gate to oversample the library at least 10-fold. Cells were collected in LB + 0.2% w/v glucose and then sorted cells were transferred into 10 mL of LB + 0.2% w/v glucose, grown at 37 °C overnight, and plasmid DNA was isolated by miniprep the next day. Three replicate MassTitr experiments were performed.

**Illumina amplicon preparation**

As described above, sorted pools were grown overnight at 37 °C in LB + 0.2% glucose and then miniprepped (Qiagen). Samples collected at a certain gate per concentration of ENAH tetramer were assigned a unique barcode/index combination. First, we PCR amplified the variable region of the library with a forward primer (Ngsfwd_1) and a corresponding reverse primer that contained one of 5 6-nucleotide (nt) index sequences for multiplexing (Ngsrev_1_i); the amplicon preparation scheme and all primers are listed in Table S5. 14 cycles of amplification were carried out with Phusion polymerase (NEB) using an annealing temperature of 66 °C. The resulting reaction was PCR purified with the Zymo Clean and Concentrate kit (Zymo Research) and eluted with 20 µL milliQ water. 200 ng of each reaction was cut with MmeI (NEB) at 37 °C for 1 hour and then the enzyme was heat inactivated at 65 °C for 20 minutes. To the 5' end of 15 µL of digested fragment we ligated a double stranded adapter with matching overhangs using T4 DNA ligase (NEB) at 25 °C for 30 minutes, followed by heat inactivation at 65 °C for 10 minutes. This adapter contained one of 24 5-nt barcodes and the standard Illumina forward primer. The expected ~200 bp band was gel purified with the Zymoclean Gel DNA recovery kit (Zymo Research) and eluted in 17 µL of MilliQ water. 3 µL of this reaction was PCR amplified for 10 cycles with primers Ngsfwd_2 and Ngsrev_2 in a 50µL reaction at an annealing temperature of 66 °C. The 5' and 3' Illumina adapter sequences and the reverse priming sequence were included in this step. The final product was PCR purified and eluted in MilliQ water. The DNA concentration of each pool was measured with the Qubit assay and samples were combined and run in one lane. In total, the multiplexed sample included 32 pools (corresponding to cells collected at each of 8 concentrations in 4 gates) x 3 replicates, as well as the pre-enriched input library, for a total of 97 samples distinguished by their barcode/index combination. At each stage of the preparation process, the quality and homogeneity of the amplicon was assessed through Sanger sequencing (Genewiz) and Bioanalyzer. The multiplexed sample was submitted for sequencing on a NextSeq500 instrument using paired end reads. Supplementary table 5 gives an overview of the procedures used to (1) label DNA from each pool of cells with a barcode/index indicating the ENAH EVH1 tetramer concentration and gate number for the sample and (2) to prepare the library for sequencing; primer sequences are listed, as well as the barcode/index key used for each sample.

**MassTitr data processing**

Sequences were demultiplexed and processed with an in-house script as described by Whitney et al. (manuscript in preparation). We used the sequencing data to determine the number of clonal cells (i.e. cells displaying the same peptide) that were found in each gate at each concentration. We calculated the clone read frequency in gate j at concentration k as $R_{ijk}/T_{jk}$, where $R_{ijk}$ is the number of raw reads for sequence i in gate j at concentration k, and $T_{jk}$ is the total sequencing reads for all clones obtained for gate j at concentration k, i.e. $T_{jk} = \sum_i R_{ijk}$. The clone read frequency was multiped by the total number of cells collected in each gate (which was recorded during the sorting) to calculate the total number of cells displaying sequence *i*, $C_{ijk}$, in gate *j* at each ENAH EVH1 tetramer concentration *k*. This can be used to compute an effective gate position using equation 1:

$$\sum_j C_{ijk} \bullet F_j / \sum_j C_{ijk} = F_{eff}^{i,k} \ (1)$$

Where $F_j$ values are the mid y-axis fluorescence values for each gate, $j$ = 1-4 (with values of 700, 1500, 2500, 4000 respectively). $F_{eff}^{i,k}$ was calculated for each clone across all ENAH tetramer concentrations. If this experiment is performed in such a way such that every peptide-expressing cell is collected and sequenced, then the clones-per-gate information can be used to extract a concentration-dependent signal-vs-concentration curve that can be fit to give an apparent dissociation constant, as done by Adams, et al. and Whitney, et al. (Adams et al., 2016; Whitney et al., manuscript in preparation). In our experiment, we collected and sequenced fewer cells, focusing on cells that displayed above-background binding signal, and obtained information about concentration-dependent binding but not complete titration curves.

Our data processing was focused on identifying a subset of well-behaved clones, of varying affinities, that we judged to be good candidates for subsequent analysis of binding features. We first removed clones for which we obtained < 100 reads across all bins at all concentrations and also clones that did not have a total cell count ($C_{ijk}$) of at least 25 cells in at least four concentrations. A sequence was assigned as an ENAH binder if (1) the clonal population of cells showed a concentration-dependent increase in binding signal in at least two of three replicate experiments, or (2) the displayed peptide contained an FP4 motif and showed concentration-dependent binding in at least one replicate. Among binders identified in this way, clones were further tagged as "high affinity" if they met additional requirements: (1) total cell count of greater than or equal to 80 cells for at least four concentrations, (2) greater than or equal to 10 cells found in binding gates at the lowest three concentrations (0.31 μM, 0.12 μM, 0.049 μM), (3) cells in more than one gate at the highest two concentrations (30 μM, 12 μM), and (4) clone found in at least two replicates. These filters are were chosen, based on benchmarking against validated binding clones from the screen, to extract a subset of well-behaved clones that we judged likely to be true binders (as proved to be true). Complete data corresponding to the MassTitr read counts per gate, for each sequence at each concentration are provided have been deposited at GEO with the accession number GSE166938.

A key providing information on which barcode/index pairs correspond to which sample as well as cell counts for each replicate are provided in Table S6.

**MassTitr sequence analysis**

The T7-pep library contains many point mutations and frameshifts, in addition to full-length human 36-mers. Consequently, for many hits we identified multiple closely related but non-identical sequences (Table S1). To analyze trends in our dataset, we collapsed hits that we judged to be variants of the same sequence into one representative sequence, leading to a total of 108 non-redundant sequences. If one of the sequence variants matched exactly with a sequence from the human proteome (UniprotKB/Swiss-Prot release 2020_01), that sequence was chosen as the representative sequence (UniProt Consortium, 2019). Otherwise, the sequence with the most combined total cell counts was chosen for analysis. To compare these sequences to sequences from the pre-enriched input library, we clustered amino-acid sequences from the input library reads using CD-HIT (Huang et al., 2010) with a sequence identity cut-off of 0.7, which effectively collapsed proteomic sequence variants into one representative sequence.

**Identification of putative biological interaction partners and GO analysis**
To identify putative biological interaction partners among our MassTitr hits we first removed peptides that mapped to a human protein but contained an unnatural FP4 or CXC motif due to frameshift or point mutations (highlighted in yellow in Table S1). Then we identified those peptides predicted to be intrinsically disordered, using an IUPred2A (Mészáros et al., 2018) cutoff of >0.4. Finally, we assessed the cytoplasmic localization of hits using cellular component terms from QuickGO (Binns et al., 2009). Two proteins from our list (NHSL1 and KIAA1522) did not have any associated terms with our search criteria. For these proteins, we manually curated subcellular localizations from the literature (Brooks et al., 2010) and the Human Protein Atlas (Thul et al., 2017). For proteins reported to be membrane bound, such as MIA3 (Reynolds et al., 2019), we confirmed that the regions we pulled out as hits were cytoplasmic as annotated in Uniprot (Uniprot Consortium, 2019). GO term enrichments were performed using PANTHER with a Fisher's Exact Test (Mi et al., 2019). GO biological process terms with an FDR < 0.05 were designated as enriched.

**Computational Rosetta modelling**

**Input structure:** An initial structure modeling bivalent binding was derived from two separate, peptide bound EVH1-domain structures. Structure 5NC7 (Barone et al., 2020) includes a short FP4-containing peptide (chain I) bound to the noncanonical site of the ENAH EVH1 domain (chain D), and the ENAH-ABI1 structure reported in this work provided a model of an $FP_8$ peptide bound at the canonical site, which for modeling was truncated to $FP_4$. These two structures were imported into PyMol and superimposed using the structural alignment function. After verifying that the RMSD for the alignment was low (≤ 0.5 Å), the ENAH EVH1 domain of 5NC7 domain was deleted. The two structures were then merged and exported from PyMol. The resulting PDB file contained an EVH1-domain bound to two independent FP4 peptides, and this structure was relaxed using the Rosetta Fast Relax protocol run under the default Rosetta energy function REF2015 (Park et al., 2016).

**Chain Bridging:** Linker geometries were modeling using RosettaScripts (Fleishman et al., 2011). The Rosetta BridgeChains mover, with the standard Rosetta energy function REF2015 with interchain centroid weights (*interchain_cen*), was used to link the noncanonical- and canonical-site-bound peptides. The insertion motif used for the protocol was "αLX", which specified that the mover should generate backbone coordinates for a loop of length α composed of amino acids derived from any part of Ramachandran space. Starting with α = 20, we tested shorter lengths of α until BridgeChains could no longer find a solution. The FP4 motif seeds were anchored in their starting positions using distance constraints generated using the CoordinateConstraintGenerator mover in *RosettaScripts* with a strength/deviation parameter of 0.25 arbitrary units. The noncanonical-site peptide required a secondary set of constraints to maintain the peptide in the observed docking position, so each of its residues was additionally constrained to lie within several angstroms of residue 45 of the EVH1 chain. The final structure, with the two motifs connected by a chain of poly-Val, was then relaxed using the Rosetta *Fast Relax* protocol run under the default Rosetta energy function REF2015. Because bivalent binding was possible in two different orientations that each preserve the polarity of both FP4 motifs as found in the 5NC7 and ENAH-ABI1 structures, this process was done twice, once for each direction.

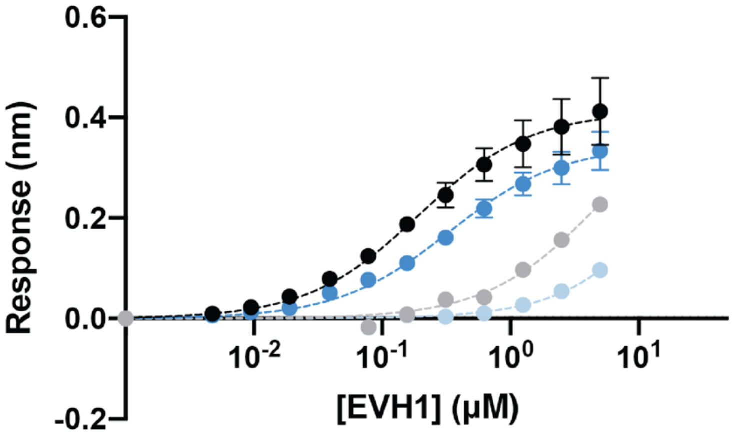


| | Sequence | $K_D$ (µM) |
|---|---|---|
| PCARE 36mer (PCARE[813-848]) | AAKSEELSCEMEGNLEHLPPPPMEVLMDKSFASLES | 0.19 ± 0.04 |
| PCARE B (PCARE[826-848]) | NLEHLPPPPMEVLMDKSFASLES | 0.32 ± 0.01 |
| PCARE C (PCARE[826-843]) | NLEHLPPPPMEVLMDKSF | n.f. |
| PCARE D (PCARE[826-840]) | NLEHLPPPPMEVLMD | n.f. |

**Figure S1. PCARE truncations.** BLI binding curves for truncations of the PCARE 36-mer. N.f. = could not be fit to give an accurate $K_D$, given this concentration range. Errors reported as the standard deviation of two replicates.

A

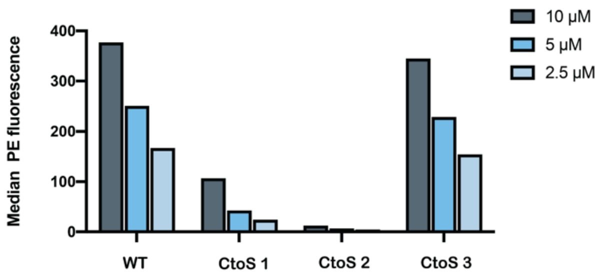


| | |
|---|---|
| OLIG3 WT | LPCPCTICQMPPPPHLSALSTANMARLSAESKDLLK |
| OLIG3 CtoS 1 | LPSPCTICQMPPPPHLSALSTANMARLSAESKDLLK |
| OLIG3 CtoS 2 | LPCPSTICQMPPPPHLSALSTANMARLSAESKDLLK |
| OLIG3 CtoS 3 | LPCPCTISQMPPPPHLSALSTANMARLSAESKDLLK |

B

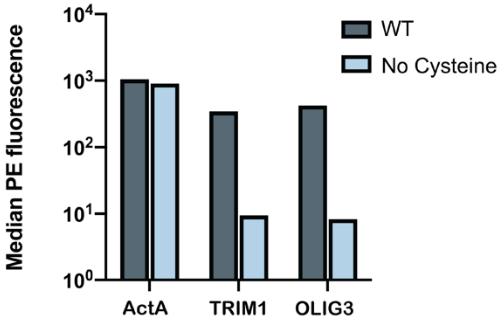


| | |
|---|---|
| ActA | GFNAPATSEPSSFEFPPPPTEDELEIIRETASSLDS |
| TRIM1 | ILSGLPAPDFIDYPERQECNCRPQESPYVSGMKTCH* |
| OLIG3 | LPCPCTICQMPPPPHLSALSTANMARLSAESKDLLK |

C

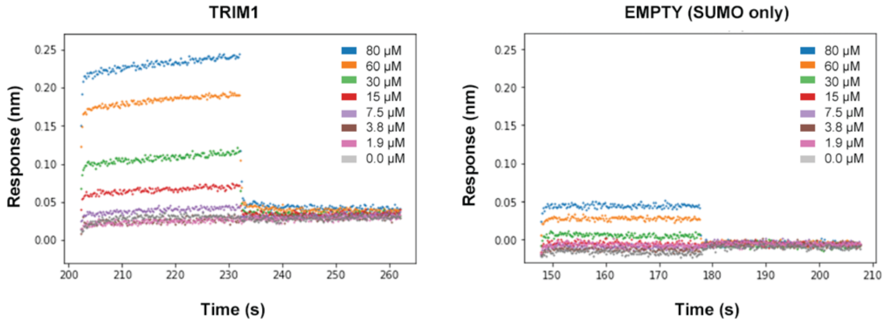


ILSGLPAPDFIDYPERQECNCRPQESPYVSGMKTCH*

**Figure S2. Peptides containing CXC motifs bind to ENAH.** 35 noncanonical (no FP4 motif) MassTitr hits contained CXC motifs. CXC motifs likely enhanced the binding affinity of weak peptides by forming disulfide bonds to the EVH1 domain in bacterial display. (A) The median fluorescence (PE) binding signal of representative CXC peptide OLIG3 mutants displayed on the surface of bacteria in the presence of either 10, 5, or 2.5 µM ENAH tetramer and 2 mM DTT. (B) The median PE binding signal of ActA peptide, or CXC-motif containing peptides TRIM1 or OLIG3, displayed on the surface of bacteria in the presence of either 5 µM ENAH tetramer, or 5 µM ENAH tetramer with no cysteines, and 2 mM DTT. (C) BLI plot for ENAH EVH1 monomer binding to a CXC-motif peptide from TRIM1 fused to SUMO (left) or to SUMO protein only (EMPTY, right). The TRIM1 binding curve did not saturate, so we did not fit a dissociation constant.

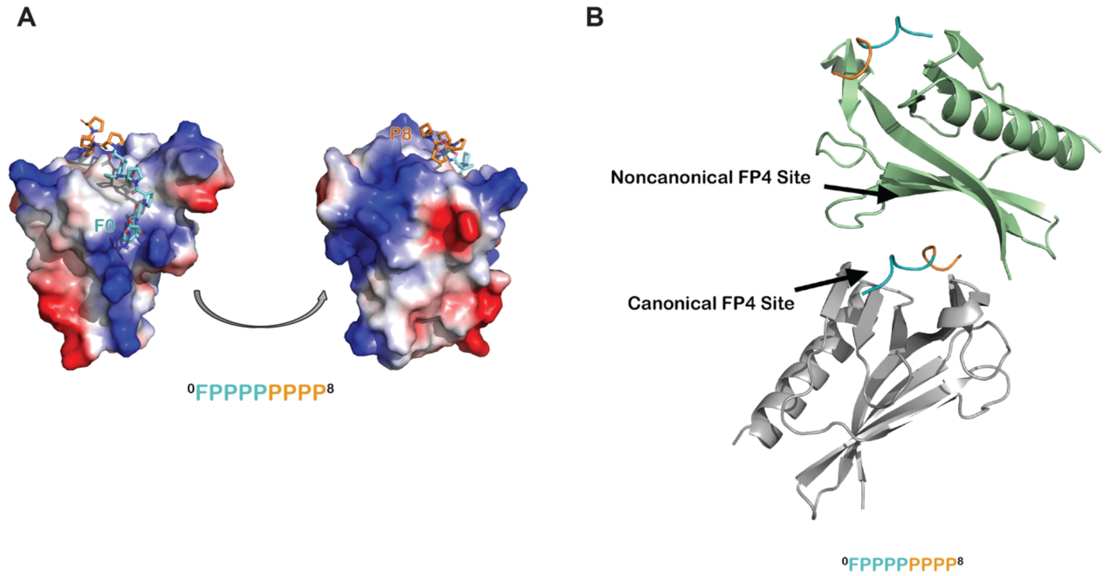


**Figure S3. Additional analyses of the ENAH EVH1-ABI1 structure.** (A) Surface representation of the ENAH EVH1 domain, colored by electrostatic potential, bound to an $FP_8$ peptide. The figure illustrates that the N- and C-terminal ends of the $FP_8$ peptide are positioned near regions of positive charge on the EVH1 domain. (B) In the ENAH EVH1-ABI1 crystal lattice, part of the $FP_8$ peptide bound to the canonical site of one EVH1 domain (gray) contacts the noncanonical site of an adjacent EVH1 domain (green). Although likely a packing artifact, this interaction mode rationalizes a possible model where two EVH1 domains can engage closely spaced FP4 motifs.

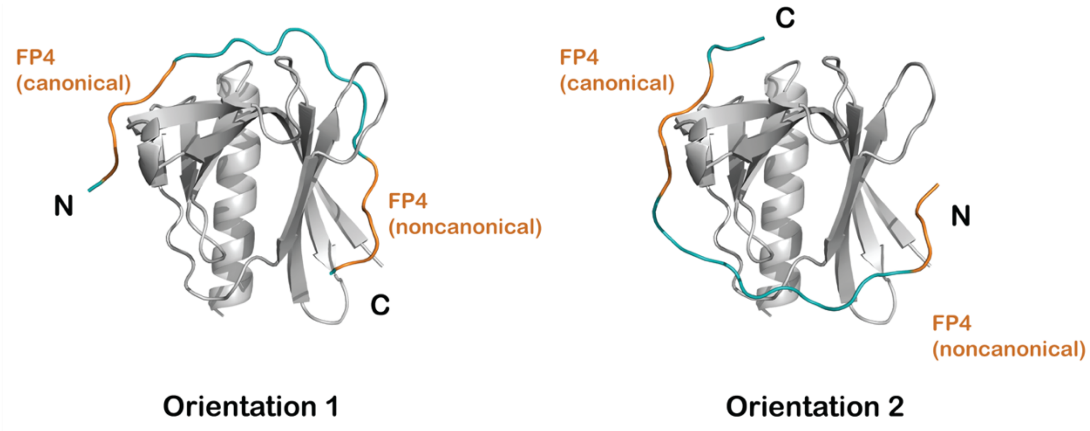


**Figure S4. Modeling bivalent binding.** (A) Models of binding in which the two FP4 motifs are linked in one of two orientations. FP4 motifs are colored in orange, and the minimum length modelled linkers (10 residues in orientation 1 and 9 residues in orientation 2) are colored blue.

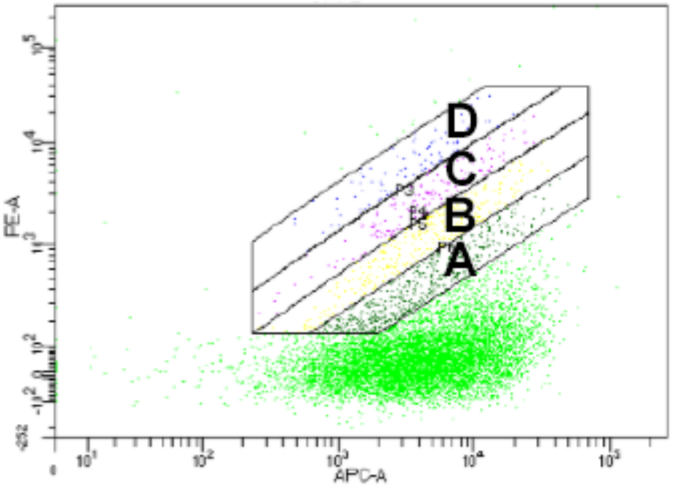


**Figure S5. Gates drawn for MassTitr FACS sort**

**Table S2. Dissociation constants for MassTitr peptides derived from human proteins binding to ENAH EVH1 domain**. In light blue and orange are peptides annotated as "high affinity" or "low affinity" by MassTitr. A peptide from ActA (gray) was used as a positive control. Affinities obtained through BLI. Error reported as the standard deviation of two replicates.

| Name | Sequence | $K_D$ (μM) [a] |
|---|---|---|
| PCARE | AAKSEELSCEMEGNLEH**LPPPP**MEVLMDKSFASLES | 0.19 ± 0.04 |
| ABI1 | FDD**FPPPP**PPPPVDYEDEEAAVVQYNDPYADGDPAW | 2.4 ± 0.1 |
| LPP | KQPGGEGDF**LPPPP**PPLDDSSALPSISGN**FPPPP**PL | 4.0 ± 3.0 |
| ZYX | ALGGA**FPPPP**PPIEES**FPPAP**LEEEI**FPSPP**PPPEE | 4.8 ± 1.3 |
| ACTA | GFNAPATSEPSSFE**FPPPP**TEDELEIIRETASSLDS | 5.2± 0.2 |
| NHSL1 | ADRSPF**LPPPP**PVTDCSQGSPLPHSPV**FPPPP**PEAL | 8.6 ± 2.0 |
| SHROOM3 | VYSMDD**FPPPP**PHTVCEAQLDSEDPEGPRPSFNKLS | 9.3 ± 0.02 |
| ABI3 | SPPPPDEELPLPLD**LPPPP**PLDGDELG**LPPPP**PGFG | 13.6 ± 1.9 |
| KIAA1522 | PPAPEEQDLSMADFPPPEEAFFSVASPEPAGPSGSP | 14.0 ± 0.3 |
| FBLIM1 | PPVLDGEDVLPDLDL**LPPPP**PPPPVLLPSEEEAPAP | 16.0 ± 0.3 |
| TNK2 | TPVVDWDARP**LPPPP**AYDDVAQDEDDFEICSINSTL | 21.3 ± 0.2 |
| TJAP | PEEELPLPAFEKLNPYPTPSPPHPLYPGRRVIEFSE | 23.9 ± 0.1 |
| FYB1 | SGSGGI**FPPPP**DDDIYDGIEEEDADDG**FPAPP**KQLD | 24.3 ± 0.7 |
| NUTM2E | VVPVMAAQVVGGTQACEGGWSQG**LPLPP**PPPPAAQL | 33.9± 1.2 |
| FHOD1 | SVPPPPP**LPPPP**PIKGP**FPPPP**PLPLAAPLPHSVPD | 50.4± 8.9 |
| SYNPO2 | KSPIAD**FPAPP**PYSAVTPPPDAFSRGVSSPIAGPAQ | 52.1± 4.9 |
| TENM1 | GSTQDVQSSPHNQFTFRP**LPPPP**PPPHACTCARKPP | 62.9± 12.5 |

**Table S3. Dissociation constants for ABI1-derived peptides binding to ENAH EVH1 domain.** Errors reported as the standard deviation of two replicates.

| Name | Sequence | $K_D$ (μM)[a] |
|---|---|---|
| ABI1 | FDD**FPPPP**PPPPVDYEDEEAAVVQYNDPYADGDPAW | 2.4 ± 0.1 |
| ABI1 b | FDD**FPPPP**PPPPVDYEDEEAAV | 3.9 ± 0.9 |
| ABI1 a | FDD**FPPPP**PPPPVDYED | 5.2 ± 0.2 |
| ABI1 ala | FDD**FPPPP**PPPPVAYAA | 8.5 ± 0.9 |
| $FDDFP_8$ | FDD**FPPPP**PPPP | 10.1 ± 0.2 |
| $FP_8$ | **FPPPP**PPPP | 28.5 ± 8.1 |
| $FP_4S_4$ | **FPPPP**SSSS | 114.4 ± 6.9 |

[a] Affinities determined by BLI as described in the methods.

**Table S4. Dissociation constants for MassTitr peptides derived from human proteins binding to EVH1 domains from VASP and EVL**. Error reported as the standard deviation of two replicates.

| Name | Sequence | VASP $K_D$ (μM)[a] | EVL $K_D$ (μM)[a] |
|---|---|---|---|
| ActA | GFNAPATSEPSSFE**FPPPP**TEDELEIIRETASSLDS | 11.2 ± 0.2 | 2.7 ± 0.3 |
| ABI1 | FDD**FPPPP**PPPPVDYEDEEAAVVQYNDPYADGDPAW | 13.6 ± 0.6 | 12.3 ± 0.4 |
| LPP | KQPGGEGDF**LPPPP**PPLDDSSALPSISGN**FPPPP**PL | 14.3 ± 0.4 | 32.3 ± 0.03 |

[a]Affinities obtained through BLI as described in the methods.

**Table S5. Comparison of affinities of single- and dual-motif peptides for ENAH WT vs. ENAH R47A.** Errors reported as the standard deviation of two replicates.

| **Name** | **Sequence** | **WT $K_D$ (μM)** [a] | **R47A $K_D$ (μM)** [a] |
|---|---|---|---|
| PCARE | AAKSEELSCEMEGNLEH**LPPPP**MEVLMDKSFASLES | .19 ± .04 | 0.33 ± .06 |
| LPP | KQPGGEGDF**LPPPP**PPLDDSSALPSISGN**FPPPP**PL | 4.0 ± 3.0 | 61.9 ± 6.1 |
| ZYX | ALGGA**FPPPP**PPIEES**FPPAP**LEEEI**FPSPP**PPPEE | 4.8 ± 1.3 | 54.5 ± 3.1 |
| ActA | GFNAPATSEPSSFE**FPPPP**TEDELEIIRETASSLDS | 5.2 ± 0.2 | 8.0 ± 0.3 |
| NHSL1 | ADRSPF**LPPPP**PVTDCSQGSPLPHSPV**FPPPP**PEAL | 8.6 ± 2.0 | 55.4 ± 1.0 |

[a] Affinities determined by BLI as described in the methods.

**Table S7. Refinement table for ENAH-ABI1 structure**

| | **ENAH-ABI1 (7LXE)** |
|---|---|
| **Wavelength** | |
| **Resolution range** | 38.74 - 1.88 (1.947 - 1.88) |
| **Space group** | C 2 2 21 |
| **Unit cell** | 56.067 77.485 66.167 90 90 90 |
| **Total reflections** | 157527 (15746) |
| **Unique reflections** | 12056 (1185) |
| **Multiplicity** | 13.1 (13.1) |
| **Completeness (%)** | 98.91 (98.08) |
| **Mean I/sigma(I)** | 15.74 (0.92) |
| **Wilson B-factor** | 47.69 |
| **R-merge** | 0.06278 (2.462) |
| **R-meas** | 0.0654 (2.561) |
| **R-pim** | 0.01806 (0.6999) |
| **CC1/2** | 0.999 (0.743) |
| **CC*** | 1 (0.923) |
| **Reflections used in refinement** | 11941 (1176) |
| **Reflections used for R-free** | 1196 (119) |
| **R-work** | 0.2447 (0.5842) |
| **R-free** | 0.2761 (0.7129) |
| **CC(work)** | 0.954 (0.735) |
| **CC(free)** | 0.964 (0.573) |
| **Number of non-hydrogen atoms** | 942 |
| **macromolecules** | 942 |
| **solvent** | |
| **Protein residues** | 122 |
| **RMS(bonds)** | 0.006 |
| **RMS(angles)** | 0.76 |

| | |
|---|---|
| **Ramachandran favored (%)** | 96.61 |
| **Ramachandran allowed (%)** | 3.39 |
| **Ramachandran outliers (%)** | 0.00 |
| **Rotamer outliers (%)** | 0.00 |
| **Clashscore** | 2.17 |
| **Average B-factor** | 75.74 |
| **macromolecules** | 75.74 |
| **solvent** | |

Statistics for the highest-resolution shell are shown in parentheses.

Note that the ABI1 $FP_8$ peptide is numbered 120-129 in accordance with its respective fusion protein numbering.

### SI References


1. Barone M, Müller M, Chiha S, Ren J, Albat D, Soicke A, Dohmen S, Klein M, Bruns J, van Dinther M, Opitz R, Lindemann P, Beerbaum M, Motzny K, Roske Y, Schmieder P, Volkmer R, Nazaré M, Heinemann U, Oschkinat H, … Kühne R. Designed nanomolar small-molecule inhibitors of Ena/VASP EVH1 interaction impair invasion and extravasation of breast cancer cells. *Proc. Natl. Acad. Sci. U.S.A*. **117**, 29684–29690 (2020).
2. Binns D, Dimmer E, Huntley R, Barrell D, O'Donovan C, Apweiler R. QuickGO: a web-based tool for Gene Ontology searching. *Bioinformatics* **25**, 3045–3046 (2009).
3. Brooks SP, Coccia M, Tang HR, Kanuga N, Machesky LM, Bailly M, Cheetham ME, Hardcastle AJ. The Nance-Horan syndrome protein encodes a functional WAVE homology domain (WHD) and is important for coordinating actin remodelling and maintaining cell morphology. *Hum. Mol. Genet.* **19,** 2421–2432 (2010).
4. Chen XJ, Squarr AJ, Stephan R, Chen B, Higgins TE, Barry DJ, Martin MC, Rosen MK, Bogdan S, Way M. Ena/VASP proteins cooperate with the WAVE complex to regulate the actin cytoskeleton. *Dev. Cell* **30**, 569–584 (2014).
5. Corral-Serrano JC, Lamers IJC, van Reeuwijk J, Duijkers L, Hoogendoorn ADM, Yildirim A, Argyrou N, Ruigrok RAA, Letteboer SJF, Butcher R, van Essen MD, Sakami S, van Beersum SEC, Palczewski K, Cheetham ME, Liu Q, Boldt K, Wolfrum U, Ueffing M, Garanto A, Roepman R, Collin RWJ. PCARE and WASF3 regulate ciliary F-actin assembly that is required for the initiation of photoreceptor outer segment disk formation. *Proc. Natl. Acad. Sci. U.S.A.* **117**, 9922–9931 (2020).
6. Drees B, Friederich E, Fradelizi J, Louvard D, Beckerle MC, Golsteyn RM. Characterization of the interaction between zyxin, and members of the Ena/vasodilator-stimulated phosphoprotein family of proteins. *J. Biol. Chem.* **275**, 22503-22511 (2000).
7. Fleishman SJ, Leaver-Fay A, Corn JE, Strauch EM, Khare SD, Koga N, Ashworth J, Murphy P, Richter F, Lemmon G, Meiler J, Baker D. RosettaScripts: a scripting language interface to the Rosetta macromolecular modeling suite. *PLOS ONE* **6**, e20161 (2011).
8. Huang Y, Niu B, Gao Y, Fu L, Li W. CD-HIT Suite: a web server for clustering and comparing biological sequences*. Bioinformatics* **26**, 680–682 (2010).
9. Krause M, Sechi AS, Konradt M, Monner D, Gertler FB, Wehland J. Fyn-binding protein (Fyb)/SLP-76-associated protein (SLAP), Ena/vasodilator-stimulated phosphoprotein (VASP) proteins and the Arp2/3 complex link T cell receptor (TCR) signaling to the actin cytoskeleton. *J. Cell Biol.* **149**, 181–194 (2000).
10. Krol A, Henle SJ, Goodrich LV. Fat3 and Ena/VASP proteins influence the emergence of asymmetric. *Development* **143**, 2172-2182 (2016).
11. Law A, Jalal S, Mosis F, Pallet T, Guni A, Brayford S, Yolland L, Marcotti S, Levitt JA, Poland SP, Rowe-Sampson M, Jandke A, Köchl R, Pula G, Ameer-Beg SM, Stramer BM, Krause M. Nance-Horan Syndrome-like 1 protein negatively regulates Scar/WAVE-Arp2/3 activity and inhibits lamellipodia stability and cell migration. bioRxiv [Preprint] (2020). https://doi.org/10.1101/2020.05.11.083030 (accessed 2 March 2020).

12. Mészáros B, Erdos G, Dosztányi Z. IUPred2A: context-dependent prediction of protein disorder as a function of redox state and protein binding. *Nucleic Acids Res.* **46**, W329–W337 (2018).
13. Mi H, Muruganujan A, Huang X, Ebert D, Mills C, Guo X, Thomas PD. Protocol Update for large-scale genome and gene function analysis with the PANTHER classification system (v.14.0). *Nat. Protoc.* **14**, 703–721 (2019).
14. Moeller MJ, Soofi A, Braun GS, Li X, Watzl C, Kriz W, Holzman LB. (2004). Protocadherin FAT1 binds Ena/VASP proteins and is necessary for actin dynamics and cell polarization. EMBO J. *23*, 3769–3779.
15. Petit MM, Fradelizi J, Golsteyn RM, Ayoubi TA, Menichi B, Louvard D, Van de Ven WJ, Friederich E. LPP, an actin cytoskeleton protein related to zyxin, harbors a nuclear export signal and transcriptional activation capacity. *Mol. Biol. Cell* **11**, 117–129 (2000).
16. Park H, Bradley P, Greisen P Jr, Liu Y, Mulligan VK, Kim DE, Baker D, DiMaio F. Simultaneous Optimization of Biomolecular Energy Functions on Features from Small Molecules and Macromolecules. *J. Chem. Theory Comput.* **12**, 6201–6212 (2016).
17. Plageman TF Jr, Chung MI, Lou M, Smith AN, Hildebrand JD, Wallingford JB, Lang RA. Pax6-dependent Shroom3 expression regulates apical constriction during lens placode invagination. *Development* **137**, 405–415 (2010).
18. Puleo JI, Parker SS, Roman MR, Watson AW, Eliato KR, Peng L, Saboda K, Roe DJ, Ros R, Gertler FB, Mouneimne G. Mechanosensing during directed cell migration requires dynamic actin polymerization at focal adhesions. *J. Cell Biol.* **218**, 4215–4235 (2019).
19. Reynolds HM, Zhang L, Tran DT, Ten Hagen KG. Tango1 coordinates the formation of endoplasmic reticulum/Golgi docking sites to mediate secretory granule formation. *J. Biol. Chem.* **294**, 19498-19510 (2019):
20. Riquelme DN, Meyer AS, Barzik M, Keating AE, Gertler FB. Selectivity in subunit composition of Ena/VASP tetramers. *Biosci. Rep.* **35**, e00246 (2015).
21. Tani K, Sato S, Sukezane T, Kojima H, Hirose H, Hanafusa H, Shishido T. Abl interactor 1 promotes tyrosine 296 phosphorylation of mammalian enabled (Mena) by c-Abl kinase. *J. Biol. Chem*. **278**, 21685–21692 (2003).
22. Thul PJ, Åkesson L, Wiking M, Mahdessian D, Geladaki A, Ait Blal H, Alm T, Asplund A, Björk L, Breckels LM, Bäckström A, Danielsson F, Fagerberg L, Fall J, Gatto L, Gnann C, Hober S, Hjelmare M, Johansson F, Lee S, Lindskog C, Mulder J, Mulvey CM, Nilsson P, Oksvold P, Rockberg J, Schutten R, Schwenk JM, Sivertsson Å, Sjöstedt E, Skogs M, Stadler C, Sullivan DP, Tegel H, Winsnes C, Zhang C, Zwahlen M, Mardinoglu A, Pontén F, von Feilitzen K, Lilley KS, Uhlén M, Lundberg E. A subcellular map of the human proteome. A subcellular map of the human proteome. *Science* **356**, eaal3321 (2017).
23. UniProt Consortium. UniProt: a worldwide hub of protein knowledge. *Nucleic Acids Res.* **47**, D506–D515 (2019).
24. van der Ven PF, Ehler E, Vakeel P, Eulitz S, Schenk JA, Milting H, Micheel B, Fürst DO. Unusual splicing events result in distinct Xin isoforms that associate differentially with filamin c and Mena/VASP. *Exp. Cell Res.* **312**, 2154–2167 (2006).

25. Zhang Y, Tu Y, Gkretsi V, Wu C. Migfilin interacts with vasodilator-stimulated phosphoprotein (VASP) and regulates VASP localization to cell-matrix adhesions and migration. *J. Biol. Chem.* **281**, 12397–12407 (2006).
